# Molecular mechanism underlying substrate recognition of the peptide macrocyclase PsnB

**DOI:** 10.1101/2021.03.09.434688

**Authors:** Inseok Song, Younghyeon Kim, Jaeseung Yu, Su Yong Go, Hong Geun Lee, Woon Ju Song, Seokhee Kim

## Abstract

Graspetides, also known as omega-ester-containing peptides (OEPs), are a family of ribosomally synthesized and post-translationally modified peptides (RiPPs) bearing side-to-side macrolactone or macrolactam linkages. Here we present molecular details of the precursor recognition of the macrocyclase enzyme PsnB in the biosynthesis of plesiocin, a Group 2 graspetide. Biochemical analysis revealed that, in contrast to other RiPPs, the core region of the plesiocin precursor peptide noticeably enhanced the enzyme-precursor interaction via the conserved glutamates. We obtained four crystal structures of symmetric or asymmetric PsnB dimers including those with a bound core peptide and a nucleotide, and suggest that the highly conserved Arg213 at the enzyme active site specifically recognizes a ring-forming acidic residue and escorts it to ATP for phosphorylation. Collectively, this study provides insights into the mechanism underlying substrate recognition in the graspetide biosynthesis, and lays a foundation for engineering new variants.

## Introduction

Natural products have been the main source of therapeutic leads owing to their structural diversity and bioactivities^1^. Ribosomally-synthesized and post-translationally modified peptides (RiPPs) are a major class of natural products that exhibit a broad scope of chemistry and diverse biological functions^2–6^. RiPPs are initially synthesized as a precursor peptide by ribosomes, and subsequently modified by enzymes that install a classdefining structural motif^2,6^. The precursor peptide is frequently composed of two segments, a leader peptide (LP), which is the site for enzyme recognition and is often cleaved during biosynthesis, and a core peptide (CP), which is the site for chemical modification. Because substrate recognition does not solely rely on the direct binding of a modification site to the enzyme active site, modular RiPP biosynthesis often leads to the natural evolution of highly diverse CPs and facile engineering of variants with higher chemical diversity.

Many structures of the enzyme-LP complex have provided molecular details of LP recognition^7–19^. However, our understanding of CP recognition and the following primary modification process has been largely limited because only a few enzyme structures with a bound CP are known. Recent examples of CP-bound structures include the radical *S*-adenosylmethionine (rSAM) enzyme CteB in ranthipeptide biosynthesis^15^, backbone *N*-methyltransferase OphMA in borosin biosynthesis^20,21^, lantipeptide dehydratase NisB^22^, and, while not an RiPP enzyme, thioamide-installing *Methanocaldococcus jannaschii* YcaO (*Mj*-YcaO)^23^. The difficulty of obtaining the native enzyme-CP structure may arise from the transient nature of the enzyme-substrate complex in enzyme reactions, and the apparent low affinity of CPs for the cognate enzyme in many RiPP families^17,24,25^. Therefore, the above examples show only a small part of the CP (CteB)^15^ or take advantage of the natural covalent linkage to the enzyme (OphMA)^20,21^, the artificial fusion of a substrate mimic to the enzyme (NisB)^22^, or the synonymous activity of the non-RiPP enzyme (*Mj*-YcaO)^23^.

Graspetides, formerly known as omega-ester-containing peptides (OEPs), are RiPPs that harbor side-to-side macrolactone or macrolactam linkages between an acidic residue (Asp or Glu) and a hydroxyl- or amine-containing residue (Ser, Thr, or Lys)^6^. The prototypic members of this RiPP family are microviridins (Group 1 graspetides) that have a cage-like tricyclic structure with two lactones and one lactam on the highly conserved TxRxPSDxDE core motif^26–28^. Recent bioinformatic and biochemical analyses revealed 11 new or putative groups of graspetides that have novel core motifs and ring connectivities, including plesiocins (Group 2 graspetides; Fig. 1a)^29–32^. The presence of topologically diverse graspetides suggests that the formation of multicyclic peptides by side-to-side crosslinking is a simple but powerful strategy to expand the chemical space of peptides both in the evolutionary pathway of nature^31,33^ and for engineering novel functionalities^34–36^.

**Figure 1.**
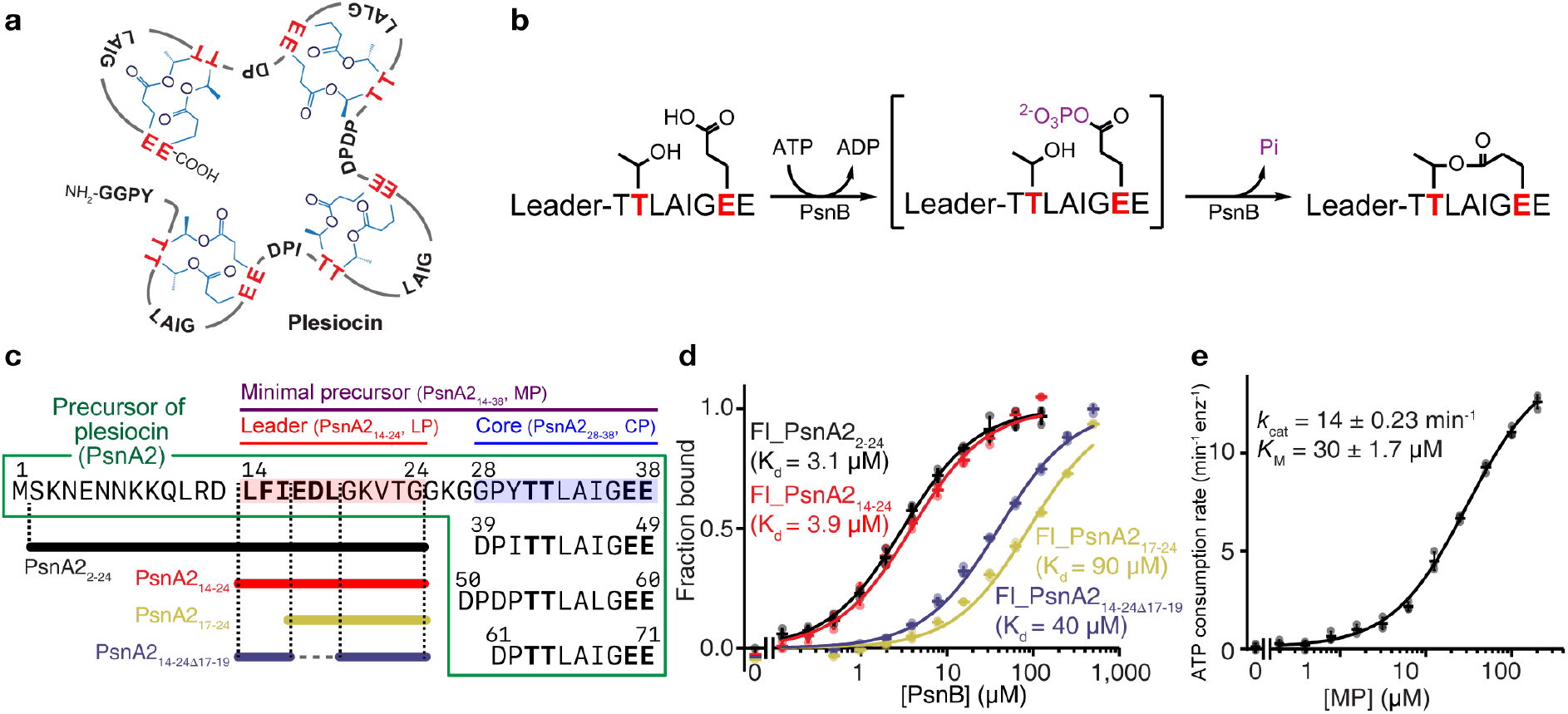
Introduction of plesiocin and its minimal precursor peptide. **a**, Structure of plesiocin, a Group 2 graspetide. Plesiocin consists of four core motifs (TTxxxxEE) and each core contains two macrolactone linkages between Thr and Glu. **b**, Suggested mechanism underlying macrolactone formation in plesiocin biosynthesis. Using ATP, ATP-grasp macrocyclase (PsnB) phosphorylates the carboxyl side-chain of glutamate in the core region of the precursor (PsnA2) and mediates nucleophilic substitution by threonine to form the macrolactone. **c**, Minimal precursor (PsnA2_14-38_, MP, overlined with purple) was designed from the native sequence of PsnA2. MP contains the key region of the leader peptide (PsnA2_14-24_, LP; red) and one core peptide (PsnA2_28-38_, CP; blue). Variants of LP used in **d** are shown as solid lines below PsnA2. **d**, The affinity of the LP variants was determined by measuring the fluorescence anisotropy of fluorophore-labeled LP variants (0.1 μM) with different concentrations of PsnB. **e**, ATP consumption by PsnB (0.4 μM) was measured with ATP (5 mM) and different concentrations of the MP. Data are presented as dot plots with mean ±1 SD (n = 3 independent experiments) and fitted to a hyperbolic equation (**d** and **e**).

ATP-grasp enzymes are responsible for the biosynthesis of macrolactone and macrolactam rings in graspetides^27,28^. Using ATP, they initially activate a carboxyl side chain by phosphorylation and mediate the nucleophilic substitution of a hydroxyl or amine group to form an ester or amide bond (Fig. 1b). Two crystal structures of the ATP-grasp enzymes in microviridin biosynthesis, the apo form of MdnB and the MdnC-LP complex, revealed a novel LP-binding domain and a distinct conformational change of the enzyme upon the leader binding—the movement of the ß9ß1O hairpin by 25 Å opens the core binding site^17^. These structures, however, showed neither the CP nor the nucleotide in the electron density maps, leaving a question about how the enzyme recognizes the CP and proceeds to the specific phosphorylation and ligation of the acidic residue.

Here, we studied the mechanism underlying precursor recognition using plesiocin biosynthesis as a model system (Fig. 1a). Biochemical studies revealed that the binding of the LP and nucleotide promotes CP binding, and that, unexpectedly, CP binding strengthens the enzyme-precursor interaction. Four crystal structures of PsnB, including those presenting a rare example of RiPP enzyme structure with well-resolved LP and CP, reveal residues involved in the binding of nucleotides, LP, or CP, and show the conformational changes of the PsnB dimer for substrate recognition. Collectively, these results suggest a molecular mechanism for the precursor binding and activation of graspetide biosynthetic enzymes.

## Results

### A minimal precursor peptide recapitulates enzyme reaction

To study the molecular details of the macrocyclization reaction in graspetides, we chose plesiocin biosynthesis as a model system. Previously, we reported that an ATP-grasp enzyme, PsnB, installs two macrolactones on each of the four conserved core motifs of the precursor, PsnA2^29^ (Figs. 1a-c). To quantitatively analyze the multiple steps of the macrocyclization reaction—precursor binding, ATP consumption, and macrolactone formation—we started by finding a minimal precursor peptide that contains only one core motif and a minimal LP. Our previous report showed that the complete macrocyclization reaction requires the conserved region of the LP, LFIEDL (PsnA2_14-19_; Supplementary Fig. 1a), but not the N-terminal 12 residues (PsnA2_1-12_)^29^. Here, we tested whether LFIEDL is required for enzyme binding by measuring fluorescence anisotropy of fluorophore-labeled LPs of various lengths (Fig. 1c). Indeed, the LFIEDL region was essential for tighter enzyme binding, whereas the N-terminal 13 residues (PsnA2_1-13_) were dispensable (Fig. 1d). Therefore, we chose PsnA2_14-38_ as the minimal precursor peptide (MP) for biochemical and structural studies.

We first measured the ATPase activity of PsnB with increasing amounts of MP. The *K*_M_ and *k_cat_* values were 30 μM and 14 min^-1^, respectively (Fig. 1e), which are similar to those of AMdnC, an enzyme for Group 1 graspetide (27 μM and 12 min^-1^)^37^. ATP titration resulted in *K_M_* and *K*_cat_ of 92 μM and 16 min^-1^ (Supplementary Fig. 1b), of which *K*_M_ is in the middle of those of non-RiPP ATP-grasp enzymes, glutathione synthetase (240 μM)^38^ and D-ala-D-ala ligase (49 μM)^39^. We also found that 10 mM Mg^2+^ was sufficient for the maximal activity of PsnB (Supplementary Fig. 1c). By monitoring water extraction in the MALDI spectrum, we observed that the modification rate of MP was similar to that of the wild-type PsnA2 (Supplementary Fig. 1d)^29^.

To obtain evidence for the suggested mechanism underlying ATP-grasp enzymes (Fig. 1b), we added hydroxylamine (NH_2_OH) to the reaction solution, which can trap the acyl-phosphate intermediate^40–42^. Although 0.5 M NH_2_OH severely reduced PsnB activity, we detected two trapped peptides: one with the NH_2_OH-added Glu37, and the other with the NH_2_OH-added Glu38 and one ester bond between T32 and E37 (Supplementary Fig. 2). These results are consistent with the activation of the carboxylate by ATP-grasp enzyme^43^ (Fig. 1b) and with the reaction order of PsnA2—the T32-E37 inner ring forms prior to the T31-E38 outer ring^29^.

### Leader peptide activates enzyme

We next investigated the role of LP in enzyme behavior. By monitoring the fluorescence anisotropy of the fluorophore-labeled CP (Fl_PsnA_228-38_ or Fl_CP) and ATP consumption of PsnB, we initially observed that the LP (PsnA2_14-24_) enhances the enzyme-CP interaction and the ATPase activity by PsnB (Supplementary Figs. 3a,b). To stably attach the LP to the enzyme, we used the leader-fused PsnB (LP_PsnB), which was shown to successfully install two macrolactones in CP^36^. The fluorescence anisotropy of the fluorophore-labeled LP (Fl_PsnA2_14-24_ or Fl_LP) indicates that the LP in LP_PsnB successfully competes for the cognate binding site in PsnB (Supplementary Fig. 3c). We then measured the fluorescence anisotropy of Fl_CP with PsnB or LP_PsnB and found that Fl_CP has a much higher affinity for LP_PsnB (*K*_d_ = 11 μM) than for PsnB *(K_d_* > 100 μM; Fig. 2a). The LP-mediated enhancement of CP affinity was also observed in the class II lantipeptide synthetase HalM2^24^. The ATPase assay showed that LP_PsnB has much higher ATPase activity, which increases with incresasing CP, whereas PsnB has minimal ATPase activity regardless of the CP (Fig. 2b). As previously reported^36^, LP_PsnB, not PsnB alone, modified CP (Fig. 2c). Collectively, the LP of PsnA2 promotes later steps of the enzyme reaction—enzyme-core interaction, ATP consumption, and macrolactone formation. These results are consistent with the enzyme activation by LP in biosynthesis of microviridins and other families of RiPPs^8,17,24,35,44–47^.

**Figure 2.**
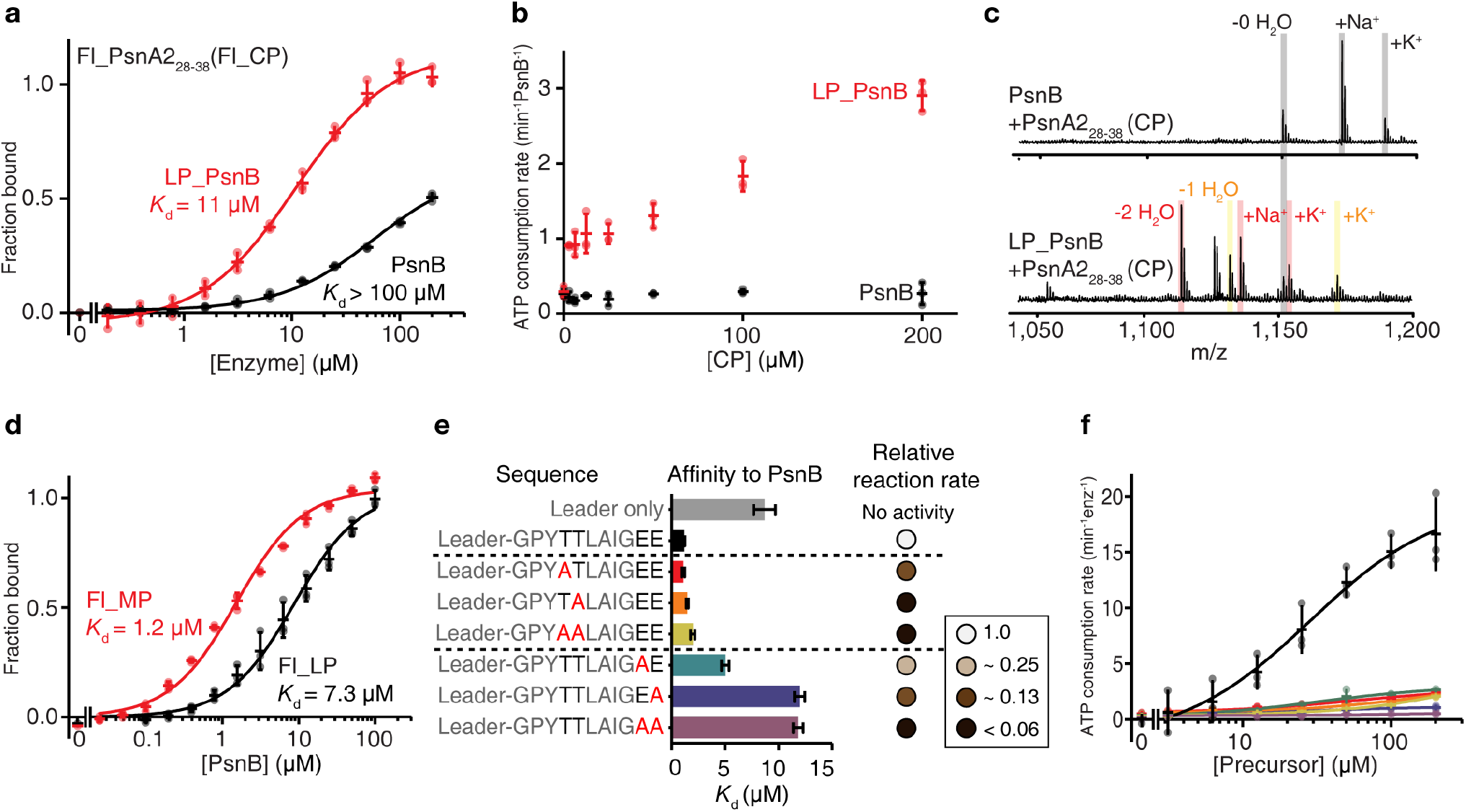
Roles of leader and core peptide in enzyme behavior. **a**, CP affinity for PsnB or the leader-fused PsnB (LP_PsnB) was determined by measuring the fluorescence anisotropy of the fluorophore-labeled CP (Fl_CP; 0.1 μM). **b**, The ATPase activity of PsnB or LP_PsnB (5 μM; 37°C) was measured with different amounts of CP. **c**, Macrocyclization of CP (200 μM) by PsnB or LP_PsnB (10 μM; 4hr, 37°C) was monitored by water extraction in MALDI spectrum. **d**, The affinity of MP or LP to PsnB was determined by fluorescence anisotropy of fluorophore-labeled MP or LP (0.1 μM). **e**, MP variants with one or two Thr-to-Ala or Glu-to-Ala mutations were tested for PsnB affinity (bar graphs) and macrocyclization reaction (circles). Affinity was determined as in **d**, and bar graphs represent the determined *K*_d_ and the error of fitting (see Supplementary Fig. 4a for data and fitting). The relative reaction rates were estimated by using MALDI spectra of the samples taken at different time points (see Supplementary Fig. 4b for MALDI spectra). **f**, The ATPase activity of PsnB with different concentrations of the MP variants shown in **e**. Data are presented as dot plots with mean ±1 SD (n = 3 independent experiments; **a, b, d,** and **f**) and fitted to a hyperbolic equation (**a, d,** and **f**).

### Core peptide enhances enzyme binding via conserved Glu

To study the effect of CP on enzyme-precursor interactions, we performed fluorescence anisotropy experiments with either Fl_LP or the fluorophore-labeled precursor (Fl_PsnA2_14-38_ or Fl_MP). Surprisingly, the precursor containing both LP and CP binds to the enzyme ~6 times tighter than the LP (Fig. 2d), suggesting that the CP of PsnA2 also contributes to the enzyme-precursor interaction. In contrast, the CP of MdnA (a microviridin precursor) did not enhance the interaction with MdnB or MdnC^17^. To determine which residues in the CP strengthen the enzyme-precursor interaction, we determined the affinity of precursor variants in which the conserved Thr or Glu residue of the CP is mutated to Ala. Ala mutations in Glu, but not in Thr, substantially reduced the affinity (4—10-fold), indicating that the conserved glutamates are critical for the affinity-enhancing effect of CP (Fig. 2e and Supplementary Fig. 4a). Interestingly, the rates of ATP consumption and macrocyclization of all these variants were considerably reduced (> 4-fold), although some of these variants still had the ring-forming Thr-Glu pair (Figs. 2e,f, and Supplementary Fig. 4b). In particular, the T31A or T32A variants that retained two conserved Glu residues also showed much lower ATPase activity than the wild-type enzyme, suggesting that Thr may help the phosphorylation of the acidic residue or the phosphorylation step may not be independent of the subsequent macrocyclization. Collectively, these results suggest that the PsnB enzyme binds to the CP primarily by the conserved Glu residues, but efficiently mediates phosphorylation and macrocyclization only with the native CP sequence, TTxxxxEE.

### Nucleotide binding enhances enzyme-core interaction

Next, we probed the effect of nucleotides on the enzyme-precursor interactions. The fluorescence anisotropy of Fl_MP or Fl_LP with ADP or AMPPNP, a nonhydrolysable analog of ATP, showed that two nucleotides enhanced the enzyme-precursor interaction by 9- and 2-folds, respectively (Fig. 3a), but not as much as the enzyme-LP interaction (Fig. 3b), indicating that nucleotides strengthen the enzyme-CP interaction. In agreement with this result, higher amounts of AMPPNP at a single concentration of PsnB (1 μM for Fl_MP and 5 μM for Fl_LP), at which only 30-40% of the peptide bound to PsnB without AMPPNP, increased the bound fraction of Fl_MP, but not of Fl_LP (Fig. 3c). These results suggest cooperative interactions among the enzyme, the precursor, including the CP, and a nucleotide. In contrast, the MdnC-MdnA interaction was not altered by the addition of ATP, ADP, or AMPPNP^17^.

**Figure 3.**
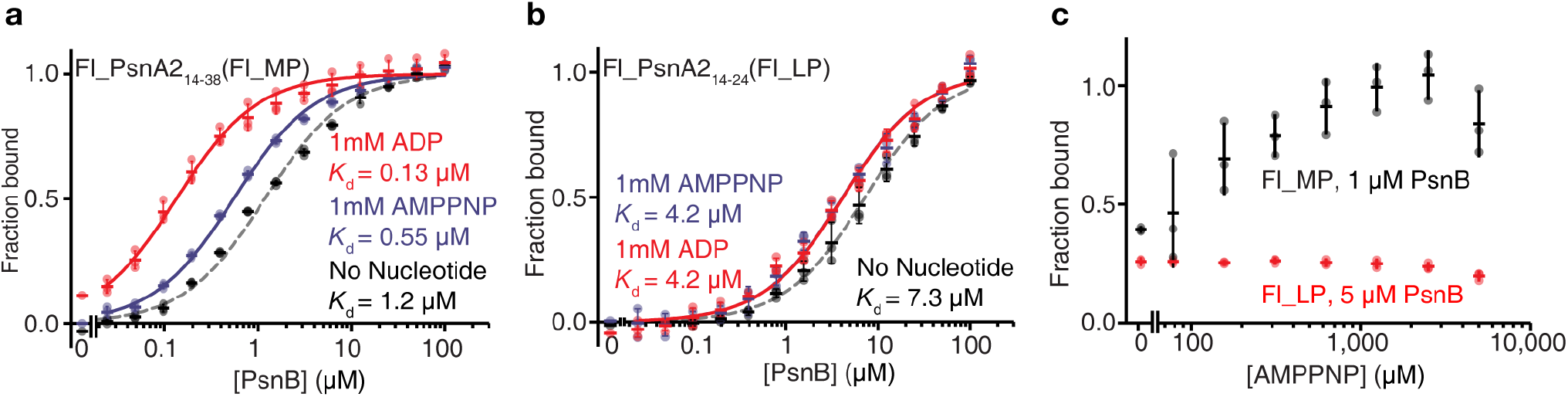
Nucleotide enhances the core affinity to PsnB. **a,b**, The affinity of MP (**a**) or LP (**b**) to PsnB in the presence of a nucleotide (1mM of ADP or AMPPNP) was determined via fluorescence anisotropy. The experiments were performed together with those in Figure 2d, and the data with no nucleotide (gray) represent those from Figure 2d. **c**, The fractions of PsnB-bound MP or LP were determined via fluorescence anisotropy with different amounts of AMPPNP. 1 μM and 5 μM of PsnB were used for MP and LP, respectively, to start with the bound fraction of 0.3-0.4 in the absence of AMPPNP. Data are presented as dot plots with mean ±1 SD (n = 3 independent experiments; **a-c**) and fitted to a hyperbolic equation (**a** and **b**).

We also determined the affinity of the precursor, the single-ring intermediate (MP-1H_2_O), and the double-ring product (MP-2H_2_O) with or without ADP (1 mM). The single-ring intermediate had the highest affinity in both conditions, although its maximal ATPase activity was approximately half that of the precursor (Supplementary Fig. 5).

### Structures of PsnB reveal core-bound asymmetric dimers

To understand the molecular mechanism underlying the enzyme-core interaction, we crystallized PnsB with a nucleotide, ADP or AMPPNP, and a minimal precursor peptide, MP or its phosphomimetic variant at Glu37 (MP(pE37)), in which the unstable acyl phosphate (CO-OPO_3_^2-^) is replaced with a more stable acyl difluorophosphonate (CO-CF_2_PO_3_^2-^; see Supplementary Fig. 6 and Supplementary Note for chemical synthesis). We determined four crystal structures containing different components (Supplementary Fig. 7a and Supplementary Table 1). Basically, the PsnB subunits show a common architecture and the conserved ATP-binding site of ATP-grasp enzymes, including MdnC and non-RiPP ATP-grasp enzymes with known structures, RimK, GshB, LysX, PurT, and DdlB (Supplementary Fig. 8)^17,48–52^. Compared with MdnC, PsnB has a longer β6α3α4 region (Arg72-Gln90) and a shorter β9β10 region (Lys172-Arg181), which corresponds to β6α3 and β9β10 in MdnC, respectively^17^. More notable differences of these PsnB structures are, however, that some PsnB subunits have well-resolved nucleotides and CP as well as LP, and that the dimers show different levels of asymmetry (Fig. 4a and Supplementary Figs. 7 and 9). Nine independent PsnB dimers are classified into four states based on their components (Fig. 4a and Supplementary Fig. 7): In two most asymmetric dimers (ENLC-E and ENCL-EN), one PsnB subunit is a full enzyme-nucleotide-leader-core complex (ENLC), while the other subunit has either a nucleotide (EN) or no additional component (E). In the other two dimers (EL-E and EL-EL) that are slightly asymmetric or almost symmetric, the PsnB subunit has either one LP (EL) or nothing (E). Although nucleotides and LPs are sometimes found in both subunits of dimers, the CP is found only in one subunit of dimers, suggesting that the binding of the CP, rather than of the LP or nucleotide, contributes more to the dimeric asymmetry. The β13β14 loop is visible only in the LP-bound PsnB subunit, but the β6α3 loop is shown only in the CP-bound PsnB subunit (Fig. 4b).

**Figure 4.**
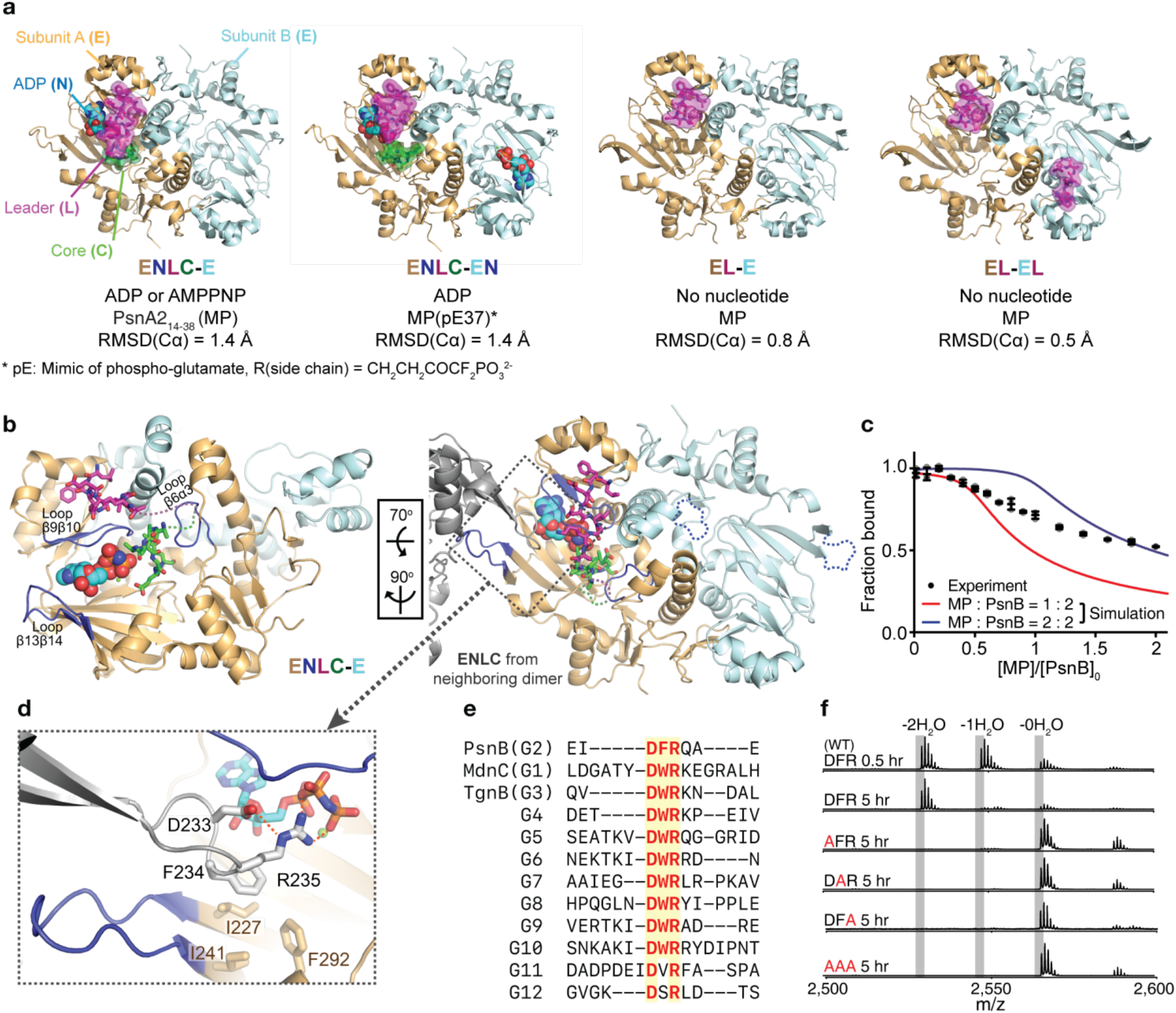
Crystal structures of PsnB reveal asymmetric dimers with bound nucleotide and core peptide. **a**, Structures of four different dimers (E, enzyme, yellow or light blue cartoons; N, nucleotide, spheres; L, leader peptide, magenta sticks/half-transparent surfaces; C, core peptide, green sticks/half-transparent surfaces). Below the structures are listed the modeled substrates and root-mean-square deviation of two superposed PsnB subunits in the dimer. **b**, Overall structure of dimeric PsnB complex (ENLC-E). Loop β6α3 and loop β13β14 are disordered in subunit B. Loop β13β14 shows intermolecular interaction with the neighboring dimer. **c**, PsnB-bound MP fractions were measured via fluorescence anisotropy of the fluorophore-labeled MP (0.1 μM) with PsnB (10 μM), ADP (1 mM), and increasing amounts of unlabeled MP. Data are presented as dot plots with mean ±1 SD (n = 3 independent experiments). The simulation curves of two models, in which the MP:PsnB stoichiometry is either 1:2 or 2:2, are shown as red and blue solid lines, respectively. **d**, The DFR residues from neighboring dimer (gray sticks/cartoon) interact with nucleotide (sky blue sticks) and hydrophobic residues (yellow sticks) from subunit A. The atoms in sticks are colored red (O), blue (N), and orange (P). The loop β13β14 and β9β10 in subunit A are colored as blue. Hydrogen-bonds are shown as red dashed lines. **e**, Sequence alignment of the DFR region in representative enzymes that synthesize 12 groups of graspetides. **f**, MALDI spectra of the reactions with PsnB variants (0.4 μM; 40 μM MP) that contain Ala mutation(s) in DFR residues.

To examine whether PsnB behaves as an asymmetric dimer in solution, we monitored the fraction of PsnB-bound Fl_MP with increasing amounts of unlabeled MP in the presence of saturating amounts of PsnB (10 μM) and ADP (1 mM). The simulation of two opposite models, in which the precursor-enzyme stoichiometry is strictly 1:2 (complete asymmetry in which only one PsnB subunit of dimer can bind to the precursor with *K*_d_ of 0.13 μM) or 2:2 (complete symmetry in which two subunits independently bind to the precursor with the same *K*_d_), shows that the bound fraction drops at approximately the precursor:enzyme ratio of 0.5 and 1, and reaches 0.25 and 0.5 at the precursor:enzyme ratio of 2, respectively (Fig. 4c). However, the experimental curve of the bound fraction decreased slightly at a ratio of 0.5, but converged to the curve of the complete symmetry model at high precursor concentrations (Fig. 4c). This result can be explained by an intermediate model in which the two subunits of the dimer can bind to the precursor with different affinities. Indeed, the original binding curves in Fig. 2d fit much better to the hill curves with Hill coefficients of 0.75 and 0.79 for MP and LP, respectively (Supplementary Fig. 10), indicating the negative cooperativity of the precursor-enzyme interaction in the PsnB dimer.

### A highly conserved DFR loop is critical for enzyme activity

Although our structures show a dimer as a fundamental unit, we found that the LP-bound PsnB subunits (ENLC and EL) also have extensive inter-dimeric interactions; the β13β14 loop, including the DFR_233-235_ residues, interacts with the β9β10 loop and a nucleotide in the neighboring dimer, or, if no nucleotide is present, occupies the nucleotide-binding site (Fig. 4d and Supplementary Fig. 11a). In particular, Arg235 interacts with the phosphates of the nucleotide, while Asp233 forms a hydrogen bond with ε-nitrogen of Arg235 and Phe234 sits on the hydrophobic surface composed of Ile227/Ile241/Phe292. Interestingly, the D(F/W)R motif is largely conserved in the ATP-grasp enzymes for graspetide biosynthesis (Fig. 4e and Supplementary Fig. 8b). RimK, GshB, and PurT also have similar sequences, but they do not bind to the nucleotide^48,49,51^. The Ala mutation on these residues resulted in the loss of macrocyclization activity, indicating that the DFR loop is critical for enzyme activity (Fig. 4f). We tested whether these inter-dimeric interactions lead to the formation of higher oligomers by using size-exclusion chromatography and fluorescence resonance energy transfer (FRET), but found no evidence for stable higher oligomers in solution (Supplementary Figs. 11b-i). Therefore, we believe that the PsnB dimers do not form higher oligomers, and the functional importance of the DFR loop can be explained by putative intramolecular interactions (Supplementary Fig. 11j), as shown in other ATP-grasp enzymes^49,53^. Alternatively, the PsnB dimers may only transiently associate together for the reaction via the DFR loop.

### Phe15 in leader peptide is important for enzyme binding

The enzyme-LP interface revealed a hydrophobic interaction between the hydrophobic pocket of PsnB (Tyr171/Ile193/Leu196/Val201/Phe203) and LP-Phe15, a putative cation-π interaction between PsnB-Arg192 and LP-Phe15, and an ionic interaction between PsnB-Arg181 and LP-Asp18 (Fig. 5a). In support of the divergence of the leader sequences, many of these residues are largely conserved only in the enzymes for Group 2a graspetide (Supplementary Figs. 8b and 12a)^31^. A similar hydrophobic interaction was also observed in the MdnC-MdnA structure (Leu196/Leu199/Phe206-Phe14)^17^. To test their contributions to the enzymeleader interaction, we characterized the leader affinity, ATPase activity, and macrocyclization activity of the Ala mutants in these residues; Leu196 and Phe203 in PsnB were critical for all these functions (4-5-fold reduction in binding affinity and >64-fold reduction in ATPase and macrocyclization activity for the Ala mutants), whereas other residues only modestly contribute (2-3-fold reduction in binding affinity and intermediate levels of ATPase and macrocyclization activity for the Ala mutants; Fig. 5b and Supplementary Figs. 12b,c,d). The affinity of the precursor variants with a mutation—F15A, D18A, or a Phe15 mutation to the pentafluorinated Phe15, F15(F5)F, which can substantially reduce the cation-π interaction^54^—indicates that Phe15 in the LP plays an important role in the enzyme-LP interaction, and that the cation-π and ionic interactions contribute only modestly (Supplementary Fig. 12e).

**Figure 5.**
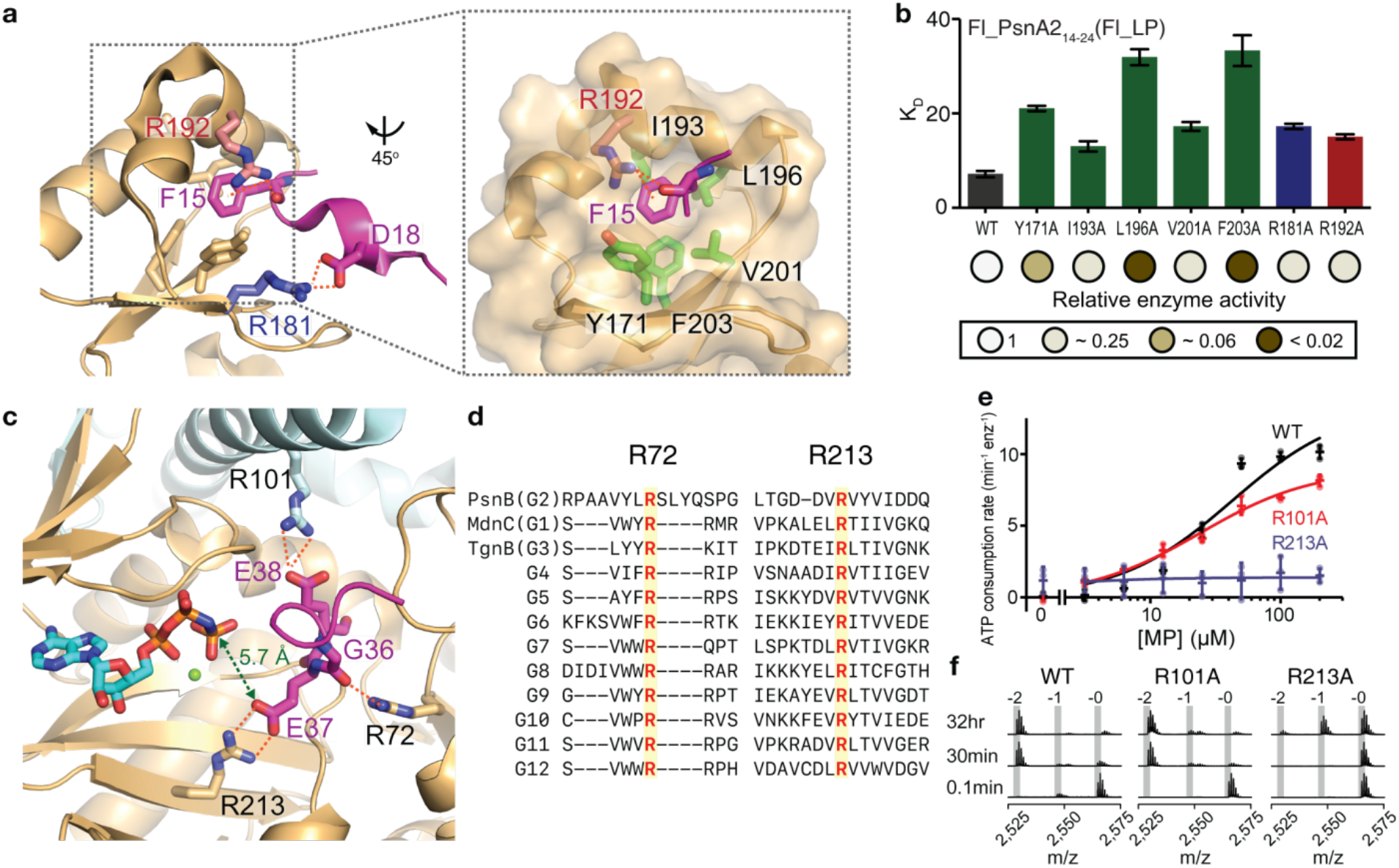
Molecular interactions between PsnB and its precursor. **a**, Interaction scheme of leader peptide (magenta sticks/cartoon) and leader-binding domain of PsnB (yellow cartoon/surface/sticks). Five hydrophobic residues of PsnB (green sticks) form a hydrophobic pocket for Phe15 of LP. Arg192 (orange sticks) of PsnB interacts with Phe15 of LP by cation-π interaction. Arg181 (blue sticks) of PsnB shows auxiliary electrostatic interaction with Asp18 of LP. **b**, The LP affinity (bar graphs) and enzymatic activity (circles) of leader-binding site mutants. Affinity was determined via fluorescence anisotropy, and bar graphs represent the determined *K*_d_ and the error of fitting (see Supplementary Fig. 12b for data and fitting). The relative activities were estimated by MALDI spectra (see Supplementary Fig. 12d for MALDI spectra). **c**, Interaction scheme of core peptide (magenta sticks/cartoon) and PsnB. Glu37 and Glu38 of CP interact with Arg213 (yellow sticks) of PsnB and Arg101 (light blue sticks) from neighboring subunit, respectively. The backbone amide of Gly36 of the core peptide interacts with Arg72 of PsnB. Carboxylate oxygen of Glu37, which forms the first ring, is located only 5.7 Å away from the phosphate of AMPPNP. **d**, Sequence alignment of the Arg72 and Arg213 regions in representative enzymes that synthesize 12 groups of graspetides. **e,f**, ATPase activity (**e**) and macrocyclization reaction (**f**) of the wild-type PsnB and its mutants carrying R101A or R213A. Data are presented as dot plots with mean ±1 SD (n = 3 independent experiments) and fitted to a hyperbolic equation (**e**).

### Conserved Arg2l3 in PsnB recognizes ring-forming Glu

Inspection of the enzyme-core interface revealed distinct ionic or hydrogen-bond interactions between the enzyme-CP residue pairs Arg213-Glu37, Arg101-Glu38, and Arg72-Gly36 (Fig. 5c). Interestingly, Arg213 interacts with the carboxyl side chain of Glu37, which is the first residue to be phosphorylated and is located near the γ-phosphate of AMPPNP (5.7 Å), and Arg101 from the neighboring subunit binds to Glu38, which forms the second ring. Arg72 and Arg213 are highly conserved in ATP-grasp enzymes for 12 groups of graspetides, whereas Arg101 is conserved only in the enzymes for Group 2a graspetides (Fig. 5d and Supplementary Fig. 8b), suggesting their different roles.

To examine the functional importance of these residues, we prepared leader-fused PsnB variants with the R72A, R101A, or R213A mutations, and found that all three Arg residues are required to tightly bind to the CP (Supplementary Fig. 13). We also purified the PsnB variants with the R101A or R213A mutation (we were unable to obtain the R72A variant), and found that Arg213 is critical for ATP consumption and macrocyclization reaction, whereas Arg101 is largely dispensable for both activities (Figs. 5e and f). These results suggest that Arg213 plays an important role in specifically recognizing the ring-forming carboxyl side chain. In agreement with this hypothesis, the crystal structure of PsnB with the phosphomimetic precursor at Glu37 revealed that Arg213 in enzyme binds to Glu38, instead of the phosphorylated Glu37 (Supplementary Fig. 9d). In this structure, we could not observe any direct interaction with the phosphorylated Glu37; therefore, it remains elusive how the hydroxyl nucleophile of Thr32 substitutes the phosphate to form an ester linkage.

### Substrate binding induces a conformational change in PsnB

Comparison of different PsnB structures revealed conformational changes in PsnB upon binding of the precursor or a nucleotide (Fig. 6a and Supplementary Video 1). Leader binding moves the α8-α9 helices of the leader binding domain to the dimeric center by 2.7 Å, generates a hydrophobic pocket to accommodate Phe15 in the LP, and relocates Arg195 out of the entry of the pocket (Supplementary Fig. 14a). Nucleotide binding moves the β9β10 loop toward the nucleotide by 1.9 Å via Lys172 and Thr180 (Supplementary Fig. 14b). The binding of both LP and CP shift the α3-α4 helices toward the active site by 2.6 Å (Figs. 6a and b). The α3-α4 helices extensively interact with the same helices in the neighboring subunit, and therefore, the movement of the rigid body of the two α3-α4 pairs appears to induce dimeric asymmetry (Fig. 6b and Supplementary Fig. 14c). The β6α3 loop becomes ordered only after the CP binds to PsnB and tightly packs the active site (Supplementary Fig. 14d). The AIGEE regions of the CPs from multiple subunit structures overlapped well with each other (Supplementary Fig. 14e). Collectively, the overall conformational change of PsnB upon the precursor and nucleotide binding establishes the space for the CP, and makes the active site compact.

**Figure 6.**
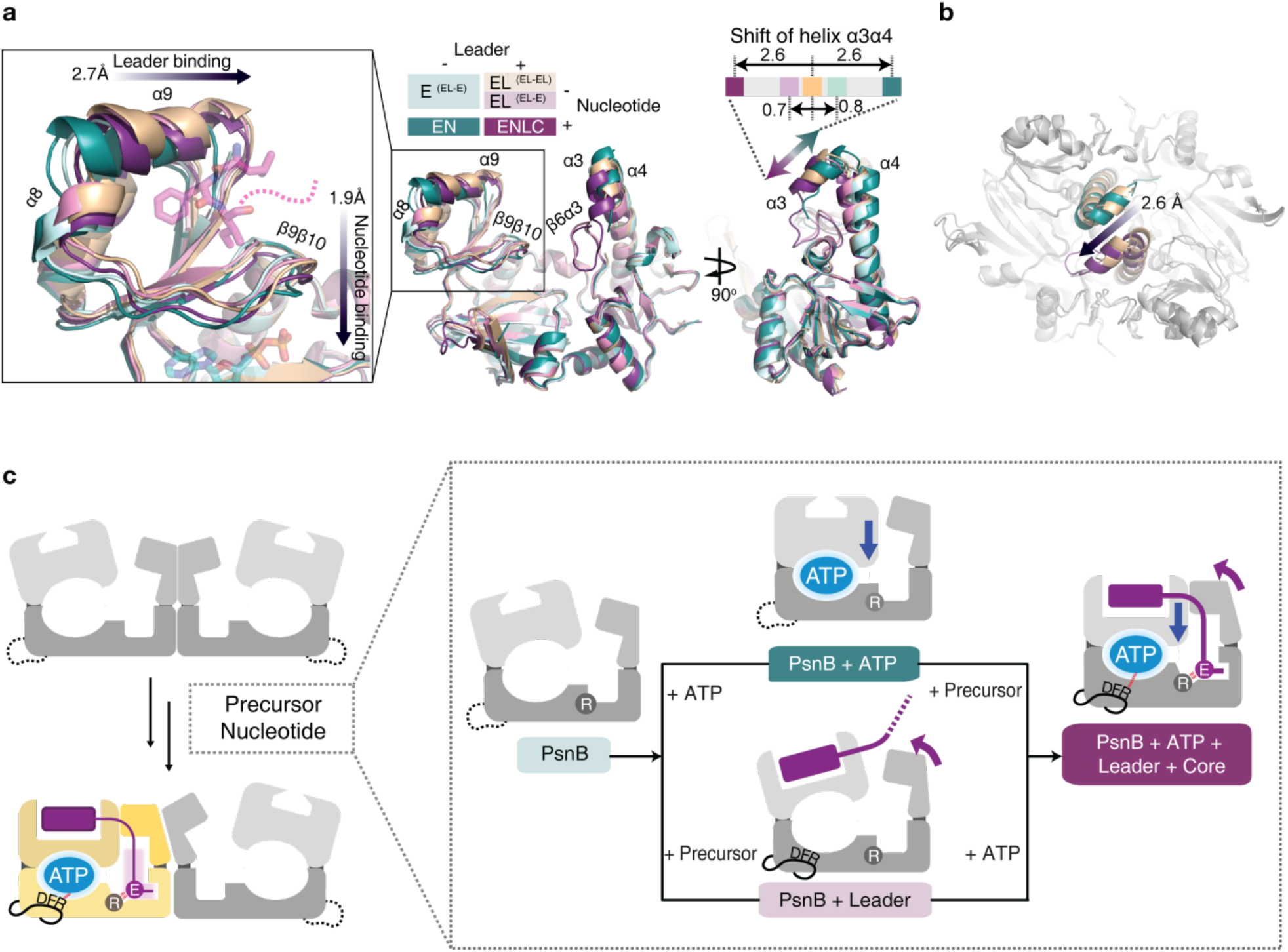
Conformational changes in PsnB dimer during substrate binding. **a**, Superposition of five PsnB subunits (E, empty PsnB, light teal; EN, enzyme-nucleotide complex, teal; EL, enzyme-leader complex, light magenta (from EL-E dimer) or orange (from EL-EL dimer); ENLC, enzyme-nucleotide-leader-core holocomplex, magenta). Leader binding moves α8α9 by 2.7 Å, nucleotide binding shifts ß9ß10 by 1.9 Å, and core binding attracts α3α4 by 2.6 Å. The β6α3 loop is ordered only when the CP is bound to PsnB. **b**, Conformational change in PsnB in the dimeric context. Two α3α4 pairs in PsnB dimer move together during the 2.6 Å shift from symmetric dimer (EL from EL-EL, orange) to asymmetric dimer (ENLC and E, magenta and teal, respectively). **c**, Suggested model for substrate recognition of PsnB. The binding of leader and ATP induces distinct conformational changes in PsnB, resulting in enhancement of the core binding. The core binding via conserved Arg213 generates the asymmetric dimer with a compact active site in core-bound PsnB.

## Discussion

By using quantitative biochemical analyses and core-bound 3D structures, we obtained a glimpse of how the ATP-grasp enzyme in plesiocin biosynthesis specifically recognizes CP, coordinates all relevant molecular interactions, and changes its conformation prior to the macrocyclization reaction. We suggest a model for substrate recognition (Fig. 6c): LP binds to PsnB mainly by hydrophobic interactions using Phe15 in the LP. PsnB also binds to ATP via several conserved residues. LP binding stabilizes the conserved DFR loop, which in turn binds to ATP. The conformational changes associated with the LP and ATP binding generate the proper space for the CP, in which the ring-forming side-chain carboxylate is specifically recognized and relocated to the site near ATP by the highly conserved Arg213 in the enzyme. The CP binding shifts the two α3-α4 pairs at the dimeric center toward the CP and stabilizes the β6α3 loop, which results in close-packing of the active site and the formation of the asymmetric dimer, in which only one subunit forms the enzyme-ATP-LP-CP holocomplex. The Arg213-bound carboxylate is then ready to move to ATP for phosphorylation. The presence of these features in enzymes for other graspetide groups remains unknown, but the high conservation of Arg213 and the D(F/W)R loop in graspetide macrocyclases indicates that their functions are conserved in graspetide biosynthesis.

The molecular details of the precursor recognition in plesiocin biosynthesis differ at several points from those in microviridin biosynthesis: (i) the CP and nucleotide also contribute to the enzyme-precursor interaction, and (ii) the enzyme forms an asymmetric dimer in an active state. The enhanced enzyme-precursor interaction by CP is most likely due to the interactions between the conserved Arg residues in the enzyme and the Glu residues in the CP. Although Arg213 is also conserved in enzymes for microviridins, this affinity-enhancing effect was not observed, suggesting the presence of another affinity-lowering factor in this enzyme. Because we observed a symmetric PsnB dimer with two LPs (EL-EL), we suggest that the reported symmetric MdnC-MdnA structure presents only one possible conformation. Collectively, plesiocin biosynthesis appears to be a better model system for understanding substrate recognition in graspetide biosynthesis.

Previously, we reported that ATP-grasp enzymes have high specificity only to the cognate core motif, and thus are not cross-reactive to CPs in other graspetide groups, which have a different core motif and ring connectivity^31^. Our structures support this idea of the evolution of high substrate specificity: (i) The core-bound structure shows a highly compact active site with well-overlapped conformations of the AIGEE region of the CPs. The β6α3 loop, which is much shorter in MdnC, is ordered only in the CP-bound PsnB, and therefore appears to generate a highly compact environment at the active site of PsnB. (ii) Arg101, which binds to Glu38 that forms the second ring, is highly conserved only in the enzymes of Group 2a graspetides. It is likely that enzymes in other groups of graspetides have different environments at their active sites to recognize their cognate core motif. Therefore, the construction of graspetide-like libraries with diverse multicyclic architectures may require diverse enzymes that produce different groups of graspetides.

There are still several unsolved questions regarding the macrocyclization reaction: what residues are involved in the phosphorylation of glutamate and the nucleophilic addition of the ring-forming Thr to generate the macrolactone linkage? Although our structure with the phosphomimetic precursor at Glu37 did not reveal residues that help the nucleophilic addition, our results suggest that two Thr residues are also involved in the phosphorylation of glutamate or that phosphorylation is coupled to nucleophilic addition. Additional structures that present snapshots of phosphorylation or nucleophilic addition may help to understand the molecular mechanisms of these steps.

Our data present a rare example of the CP-bound structure of enzymes involved in RiPP biosynthesis. We believe that this is possible because of the well-defined minimal precursor peptide and the contribution of CP to enzyme binding. Because the minimal precursor peptide, PsnA2_14-38_, is short enough for solid-phase peptide synthesis, we believe that chemical synthesis of intermediate mimics as well as mutant enzymes that may stabilize the enzyme-intermediate complex may help further dissect the macrocyclization process. The molecular details of the macrocyclization reaction will not only improve our understanding of graspetide biosynthesis, but also help the construction of graspetide-like peptide libraries to explore diverse biological functions for therapeutics and biotechnology.

## Supporting information

Supplementary Video 1.mov

## Methods

### General materials and methods

Reagents for cloning were purchased from Enzynomics (Korea) or Toyobo (Japan). Oligonucleotides were purchased from Macrogen (Korea). *Escherichia Coli* DH10β and BL21(DE3) strains were used for cloning and protein overexpression, respectively. Protein and peptide concentrations were determined by UV absorbance at 280 nm. Amino acids, coupling reagents, and resins for peptide synthesis were purchased from GL-Biochem (China). For protein purification, Ni sepharose 6 FastFlow beads were obtained from GE Healthcare (USA). Anion exchange Chromatography was performed using ÄKTA pure system (GE Healthcare) on a MonoQ 5/50 GL column (GE Healthcare). Sample purification and analysis with HPLC were performed using Agilent 1260 Infinity (Agilent, USA) on a ZORBAX SB-C18 Semi-preparative column (9.4 x 250 mm, 5 μm particle size; Agilent) and ZORBAX SB-C18 Analytical column (4.6 x 250 mm, 5 μm particle size; Agilent), respectively. 0.05% trifluoroacetic acid (TFA) in H_2_O (v/v, Solvent A) and 0.05% TFA in acetonitrile (CH_3_CN; v/v, Solvent B) were used as mobile phase of HPLC. Mass analysis was performed by Ultraflextreme MALDI-TOF/TOF mass spectrometer (Bruker Daltonics, USA).

### Cloning, overexpression and purification of PsnB variants

Plasmids and oligonucleotides used in this study are listed in Supplementary Table 2 and 3, respectively. Plasmids expressing PsnB variants were constructed using an inverse PCR method^55^. PsnB and its variants were expressed and purified as previously described^36^.

### Peptide synthesis

Minimal precursor (PsnA2_14-38_), leader peptide (PsnA2_14-24_), core peptide (PsnA2_28-38_), and its variants were synthesized by solid-phase peptide synthesis. Wang resin (200 μmol, 208 mg) was washed and soaked with 5mL of 1:1 dimethylformamide (DMF) and dichloromethane (DCM) for 20 min in a reaction vessel for resin swelling. Attachment of 1^st^ amino acid was performed by N,N-diisopropylcarbodiimide (DIC) coupling. The solution of Fmoc-AA-OH (5 equiv.), 4-dimethylaminopyridine (DMAP; 0.1 equiv.), and DIC (4 equiv.) in 2~3ml DMF was added to the reaction vessel carrying the resin, and incubated at room temperature for 2~3 days. Resins were washed three times with DMF and DCM, and were capped by acetic anhydride: acetic anhydride (10 equiv.) and DMAP (0.1 equiv) in 2 ml DCM was incubated with resins for 1 hr. After the capping, resins were washed with DMF and DCM again, and mixed with 20% (v/v) piperidine in DMF for 30 min to cleave Fmoc. After DMF and DCM wash, the 1^st^ AA-bound resins went through the sequential coupling-capping-deprotection. Coupling was performed by incubating the resins with reaction solution that contains Fmoc-AA-OH (5 equiv.), HATU/N,N-Diisopropylethylamine (DIEA; 10/5 equiv.) dissolved in DMF. Then, resins were treated with capping solution (DMF/acetic anhydride/DIEA in a 9:1:0.05 ratio) for 7 min. For the deprotection of Fmoc, resins were mixed with 20% piperidine in DMF for 30 min. Resins were washed with DMF and DCM three times between each step. All the remaining amino acids were sequentially attached by repeating coupling, capping, and deprotection. The completion of each step was monitored by tests with ninhydrine or 2,4,6-Trinitrobenzenesulfonic acid (TNBS). After the completion of the synthesis, peptides were detached from the resins by incubating the resins with cleavage cocktail (95% TFA, 2.5% deionized water, 2.5% triisopropyl silane) for 3 hrs. Cleavage solutions were then evaporated and mixed with 10-fold excess of ether:hexane solution (1:1) for peptide precipitation. The precipitated peptides were collected by centrifugation at 4,000 x g for 15min, dried, dissolved in DMSO, purified by HPLC, and lyophilized. For the synthesis of fluorophore-labeled peptides, 5(6)-Carboxyfluorescein (Acros) and serine (as a linker) were attached to the N-terminus of peptides. MS profiles of synthesized peptides are provided in Supplementary Table 4.

### Chemical synthesis of the phosphomimetic glutamate

The phosphomimetic glutamate (pGlu or pE) contains acyl difluoro-phosphonate (CO-CF2PO_3_^2-^) instead of acyl phosphate (CO-OPO_3_^2-^). Synthesis of the phosphomimic of glutamate is described in Supplementary Fig. 6: briefly, benzyl lithiodifluoromethylphosphonate was added to Fmoc-protected glutamic anhydride to furnish phosphomimetic Glu^56^. Peptide synthesis with the phosphomimetic Glu was performed in the same fashion as normal peptides.

### Fluorescence anisotropy

Fluorescence anisotropy was performed as follows: a dye-labeled peptide (0.1 μM) and 2-fold serial dilutions of PsnB variants (starting at 100-500 μM) were mixed in a buffer containing Tris-HCl (100 mM; pH 7.3), MgCl_2_ (10 mM), and KCl (50 mM) at 37°C. Fluorescence (excitation, 485 nm; emission, 535 nm, parallel or perpendicular) was monitored after 1 min incubation by using Infinite F200pro (Tecan).

### ATPase assay

ATPase assays were performed as previously described^57^. Briefly, precursor peptide variants (0-200 μM) and PsnB variants (single concentrations at 0.4-2 μM) were co-incubated at 37°C in a buffer containing ATP (5 mM; Sigma-Aldrich), NADH (400 μM; Sigma-Aldrich), pyruvate kinase (20 unit/ml; Sigma-Aldrich), L-lactic Dehydrogenase (20 unit/ml; Sigma-Aldrich), phosphoenolpyruvate (3 mM; Sigma-Aldrich), Tris-HCl (100 mM; pH 7.3), MgCl_2_ (10 mM), and KCl (50 mM). 47.5 μl of reaction mixtures without ATP were transferred into a 384 well microplate (SPL life science). 2.5 μl of ATP (100 mM) were supplied just before the measurement. After ATP were added, absorbance at 340 nm were measured using Infinite M200pro (Tecan) equipped with i-Control software. The rate of ATP hydrolysis was calculated from the following equation.

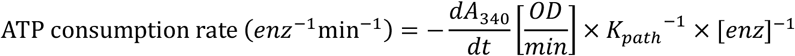

Where *K*_path_ is the molar absorption coefficient for NADH for a given optical path length. *K*_path_ is equal to 6.67×10^2^ for 50 μl well fill volume. The rates were normally corrected for background NADH decomposition of controls containing neither enzyme nor peptide.

### *In vitro* reaction and analysis of PsnA2 variants

*In vitro* reactions were performed at 37°C in solutions containing precursor peptide vaiants (40-100 μM), PsnB variants (0.5-5 μM), ATP (5 mM), Tris-HCl (100 mM; pH 7.3), MgCl_2_ (10 mM), and KCl (50 mM). At the designated time points, reaction solutions were quenched by adding the same volume of 10% (v/v) TFA. After quenching, peptides were desalted using C18 zip-tip (Millipore) and analyzed by MALDI-MS. The MS profiles are provided in Supplementary Table 4.

### Trapping the acyl-phosphate intermediate

Reaction condition was same as that of *in vitro* reaction condition except for the addition of hydroxylamine (0.5 M, final concentration) at the start of the reaction. Reaction was quenched after 4 hr incubation. Hydroxylamine-trapped peptides were monitored by MALDI, purified by HPLC and further analyzed by MALDI-MS/MS. The MS/MS profiles are provided in Supplementary Table 4. Methanolysis to confirm the connectivity of the hydroxylamine-trapped peptides with one ester bond (2-c) was performed as previously described^29^.

### Crystallization, data collection and crystallographic analysis

For PsnB-MP-ADP (7DRM), PsnB-MP-AMPPNP (7DRN), and PsnB-MP(pE37)-ADP (7DRP), purified PsnB (3.8 mg/ml) was mixed with nucleotide (5 mM), MgCl_2_ (5 mM), and 1.2 equiv. of the minimal precursor (PsnA2_14-38_) or phosphomimetic precursor (PsnA2_14-38_(pE37)). Initial screening was performed in a vapor diffusion and sitting drop format. Small crystals were identified in a reservoir condition of 0.1M sodium cacodylate pH 6.5, 0.1 M calcium acetate, and 12% PEG8000. pH, salt, and precipitant optimization were performed in hanging drop screens. Final reservoir conditions are as follows: PsnB-MP-ADP, 0.1M Na acetate pH 5.4 and 7% PEG3350; PsnB-MP-AMPPNP, 0.1M Na acetate pH 5.2 and 6% PEG3350; PsnB-MP(pE37)-ADP, 0.1M Na acetate pH 5.0 and 3% PEG3350. Crystals of PsnB complex were grown at 20°C. For PsnB-MP (7DRO), purified PsnB (7.6 mg/ml) was mixed with ADP (1 mM), MgCl_2_ (1 mM), and 1.2 equiv. of MP (PsnA2_14-38_). Initial screening was performed in the same format and small crystals were identified in a reservoir condition of 0.2 M potassium citrate tribasic monohydrate and 20 % w/v PEG3350. Same optimization processes were done to obtain optimized protein crystals. Final reservoir condition is 10% tacsimate pH 7.6 and 14% PEG3350. Crystals were frozen in liq. N2 and diffraction data were collected on beamline 7A of the Pohang Accelerator Laboratory (PAL) at a wavelength of 0.97934 Å. Data were processed by using HKL2000^58^. Molecular replacement was carried out using Phaser^59^ whereby the structure of MdnC (PDB 5IG9) was used as a model. Refinement with Phenix^60^ along with manual model rebuilding and ligand placement with COOT^61^ produced the final models. Structural figures were prepared with PyMol (https://pymol.org/2/).

### Binding stoichiometry

Binding stoichiometry between PsnB and MP was measured by fluorescence anisotropy of fluorophore-labeled MP (Fl_MP) in the same condition as previously described: 0.1 μM Fl_MP, 10 μM PsnB, and 1 mM ADP were co-incubated with 0-20 μM of non-labeled MP. Binding fraction of Fl_MP was measured with the different concentrations of non-labeled MP. Totally symmetric and totally asymmetric binding model are simulated with the *K*_d_ of 0.13 μM.

### Size-exclusion chromatography

Size exclusion chromatography was performed using ÄKTA pure system (GE Healthcare) on a Superdex 200 column (GE Healthcare). A solution of 20 mM Tris-Cl pH 8.0 and 100 mM NaCl was used as an eluent. 10 mM MgCl_2_ or 1mM ATP were used for different eluent conditions.

### Dye-labeling and fluorescence resonance energy transfer (FRET)

Purified PsnB was incubated with 10 mM Tris and 10 mM dithiothreitol (DTT) for 1 hr and then purified with gel filtration using fresh 10 mM Tris buffer. At first, labeling with phenylmaleimide (PM) was performed to monitor which cysteine residues of PsnB are preferentially labeled. Reduced PsnB (50-100 μM) and 5 equiv. phenylmaleimide were mixed in 10 mM Tris pH 8.0. After 2 hr incubation at room temperature, reaction was quenched with excess DTT. Labeled PsnB was cleaved by adding trypsin (trypsin/PsnB 1:100, *w/w)* and analyzed by MALDI-MS/MS. Alexa Flour 555 and Alexa Flour 647 (Thermo Fisher Scientific, USA) were covalently linked to native Cys residues of PsnB in the same condition of PM labeling. After dye labeling, dye-labeled PsnB was purified by sizeexclusion chromatography. FRET assays for enzyme assembly were performed with 0.2 μM donor- and acceptor-labeled PsnB using a micro-plate reader (excitation, 520 nm; emission, 570 or 670 nm; Infinite F200pro, Tecan).

## Data availability

Coordinates and structure factors for the reported crystal structures in this work were deposited in the RCSB Protein Data Bank under accession numbers 7DRM (MP- and ADP-bound PsnB), 7DRN (MP- and AMPPNP-bound PsnB), 7DRO (MP(pE37)- and ADP-bound PsnB), and 7DRP (MP-bound PsnB). Source data are provided with this paper.

## Acknowledgements

We thank Hyunbin Lee, Ga-eul Eom, Chanwoo Lee, Heejin Roh, Hyunjin Cho, and Hyojin Park for helpful discussions and technical assistance; Hyunho Chung, Yu Jin Lee, and Tae Young Im for assistance in synthesizing compounds; Yurie T Kim, Woo Jae Jeong, and Yuri Choi for advice in crystallization and crystallographic analysis; Hyeonuk Woo for help in loop modeling. We also thank the staffs of the 5C and 7A beamlines at the Pohang Light Source. This research was supported by Basic Science Research Program through the National Research Foundation of Korea (NRF) funded by the Ministry of Education (NRF-2020R1F1A1054191).

## Author contributions

I.S. and S.K. conceived the project, designed experiments, analyzed data, and wrote the paper; S.K. supervised the project; I.S. performed majority of experiments; Y.K. performed a part of mutant analysis; I.S. and J.Y. conducted crystallographic experiments under the supervision of W.J.S. and S.K.; I.S. and S.Y.G synthesized the phosphomimetic glutamate under the supervision of H.G.L. and S.K.

---

**Supplementary Video 1 | Morph of PsnB complex.** Morph between the symmetric conformation of the PsnB dimer (EL-EL) and the asymmetric PsnB dimer (ENLC-E). Precursor peptides and nucleotides are shown magenta (ribbon and stick) and cyan (stick).

