## Supplementary Information.pdf for "Molecular mechanism underlying substrate recognition of the peptide macrocyclase PsnB"

### Table of contents

**Supplementary Figures……………………………………………………………………………….S3**

Supplementary figure 1. Designed minimal precursor recapitulates the reactivity of PsnB. ……….………….S3

Supplementary figure 2. Acyl-phosphate intermediates were trapped by hydroxylamine ……………..……….S4

Supplementary figure 3. Binding of the leader peptide activates PsnB.…………………………….……………S5

Supplementary figure 4. Binding and modification property of MP variants …….…………………………………S6

Supplementary figure 5. Binding and modification of ring-containing precursors ……..…………..…….……….S7

Supplementary figure 6. Chemical synthesis scheme of a phosphomimetic glutamate…………..……………..S8

Supplementary figure 7. Four crystal structures of PsnB complexes ………………………….…...……………..S9

Supplementary figure 8. Conservation pattern of residues implicated in substrate interaction of ATP-grasp enzymes ……….……………..……………………..…………………………………………………..S10-S12

Supplementary figure 9. Electron density maps for nucleotide, LP, and CP …………………….…………..S13

Supplementary figure 10. Binding of MP and LP to PsnB shows negative cooperativity ……………….…..S14

Supplementary figure 11. DFR residues are critical for enzyme activity …………………..……….…... S15-S16

Supplementary figure 12. Characterization of leader-binding site mutants of PsnB …………….………..…..S17

Supplementary figure 13. Binding property of core-binding site mutants of leader-fused PsnB…………..…..S18

Supplementary figure 14. Precursor and nucleotide binding induces conformational change of PsnB ...……..S19

**Supplementary Tables……………………………………………………………………………S20**

Supplementary table 1. Data collection and refinement statistics for 4 structure models……..……………….S20

Supplementary table 2. Plasmid informatics used in this study……….……………………………….……….S21

Supplementary table 3. Oligonucleotides for cloning in this study….…………………….…………...….S22

Supplementary table 4. Observed and calculated mass values of MALDI data……...………...……..….S23-S25

**Supplementary Note……………………………………………………………………………S26**

**Supplementary References……………………………………………….…………………..……S29**

### Supplementary Figures

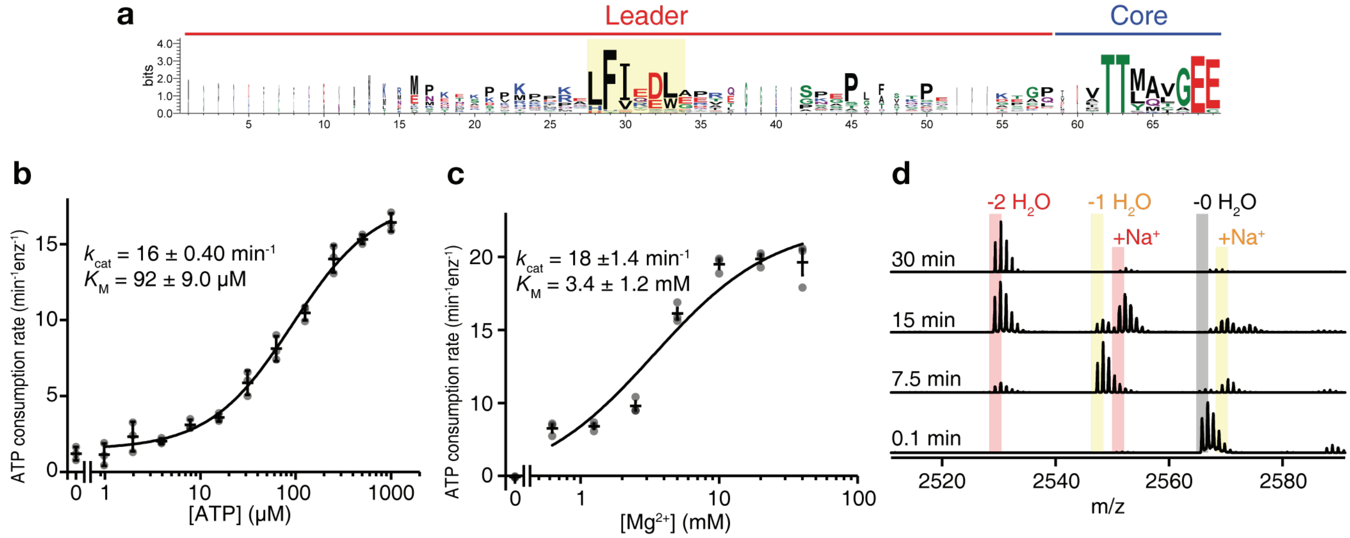


**Supplementary figure 1. Designed minimal precursor recapitulates the reactivity of PsnB. a**, Sequence logo of precursors of Group 2a graspetides including the leader peptide (red region) and one core motif (blue region). The LFIEDL region is highly conserved in Group 2a graspetides (yellow box). **b**,**c**, ATPase assays titrating with ATP (**b**) in solutions containing 0.4 μM PsnB, 200 μM minimal precursor (MP), 100 mM Tris pH 7.3, 50 mM KCl, and 10 mM $MgCl_2$, or with $Mg^{2+}$ (**c**) in solutions containing 0.4 μM PsnB, 200 μM MP, 100 mM Tris pH 7.3, 50 mM KCl, and 5 mM ATP. Data are presented as dot plots with mean ±1 SD (n = 3 independent experiments) and fitted to a hyperbolic equation. **d**, Minimal precursor was successfully modified by PsnB. 0.5 μM PsnB and 50 μM MP were co-incubated in buffer A (100 mM Tris pH 7.3, 50 mM KCl, 5 mM ATP, and 10mM $MgCl_2$) at 37°C, and the reaction solutions at designated time points were analyzed by MALDI-MS.

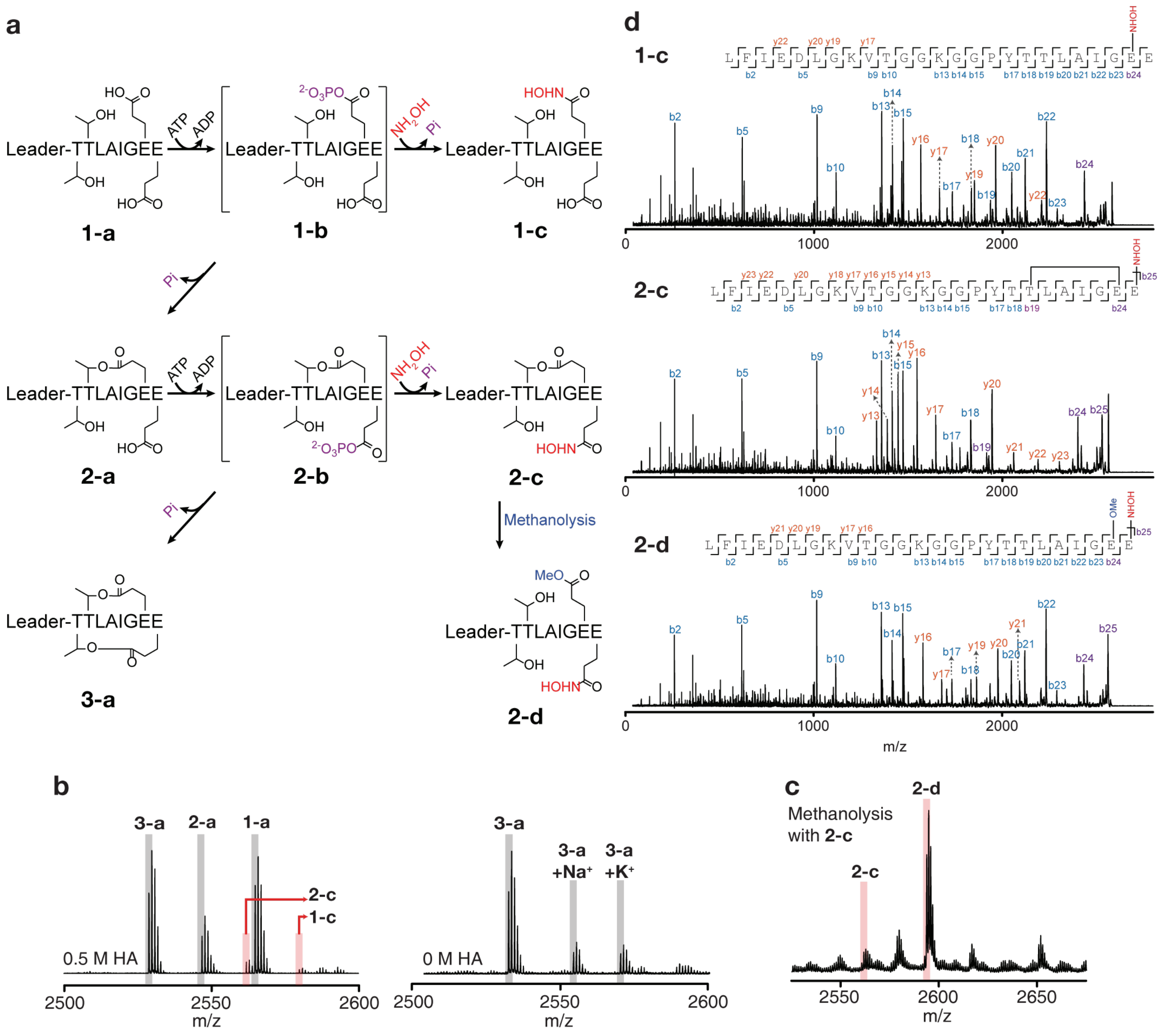


**Supplementary figure 2. Acyl-phosphate intermediates were trapped by hydroxylamine. a**, Scheme of the acyl-phosphate trapping during the macrocyclization reaction. Nucleophilic attack of hydroxylamine rather than the core threonine generates the hydroxylamine adducts of the precursor. **b**, MALDI analysis of the reaction solution with (left) or without (right) hydroxylamine. 0.5 M hydroxylamine was added to the reaction mixture containing 100 μM MP (**1-a**) and 6 μM PsnB in buffer A. The reaction mixtures were analyzed by MALDI after 4 hr incubation at 37°C. Co-incubation of 0.5 M hydroxylamine generated $NH_2OH$-added precursor peptides (**1-c** and **2-c**), which are the result of nucleophilic attack of hydroxylamine to acyl-phosphate intermediates (**1-b** and **2-b**). **c**, **2-c** was purified by HPLC and methanolysis was performed with purified **2-c** as previously reported[1]. The methanolysis product (**2-d**) was detected by MALDI. **d**, MALDI-MS/MS analysis of three hydroxylamine adducts. The connectivities of ester bonds were determined by MS/MS analysis with $NH_2OH$-added precursor peptides (**1-c** and **2-c**) and methanolysis product (**2-d**).

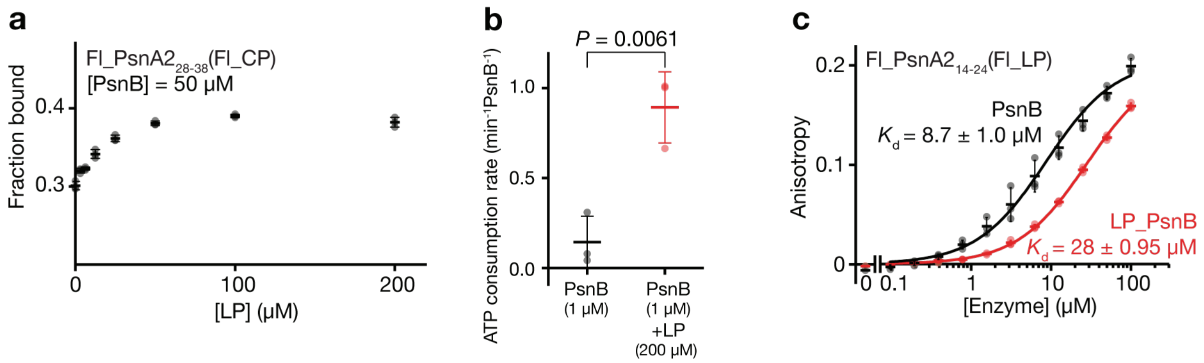


**Supplementary figure 3. Binding of the leader peptide activates PsnB.** **a**, LP enhances the CP affinity to PsnB. Fraction bound of Fl_CP (0.1 μM) to PsnB (50 μM) were determined by fluorescence anisotropy. **b**, LP enhances the ATPase activity of PsnB about 6-fold. Basal ATP consumption rate of PsnB was 0.14 $min^{-1}enz^{-1}$, whereas the addition of 200 μM LP increased the ATP consumption rate to 0.89 $min^{-1}enz^{-1}$. $P$ value < 0.01 by Student's t-test. **c**, Fluorescence anisotropy of Fl_LP (0.1 μM) to wild-type PsnB or leader-fused PsnB (LP_PsnB). Data are presented as dot plots with mean ±1 SD (n = 3 independent experiments; **a-c**) and fitted to a hyperbolic equation (**c**).

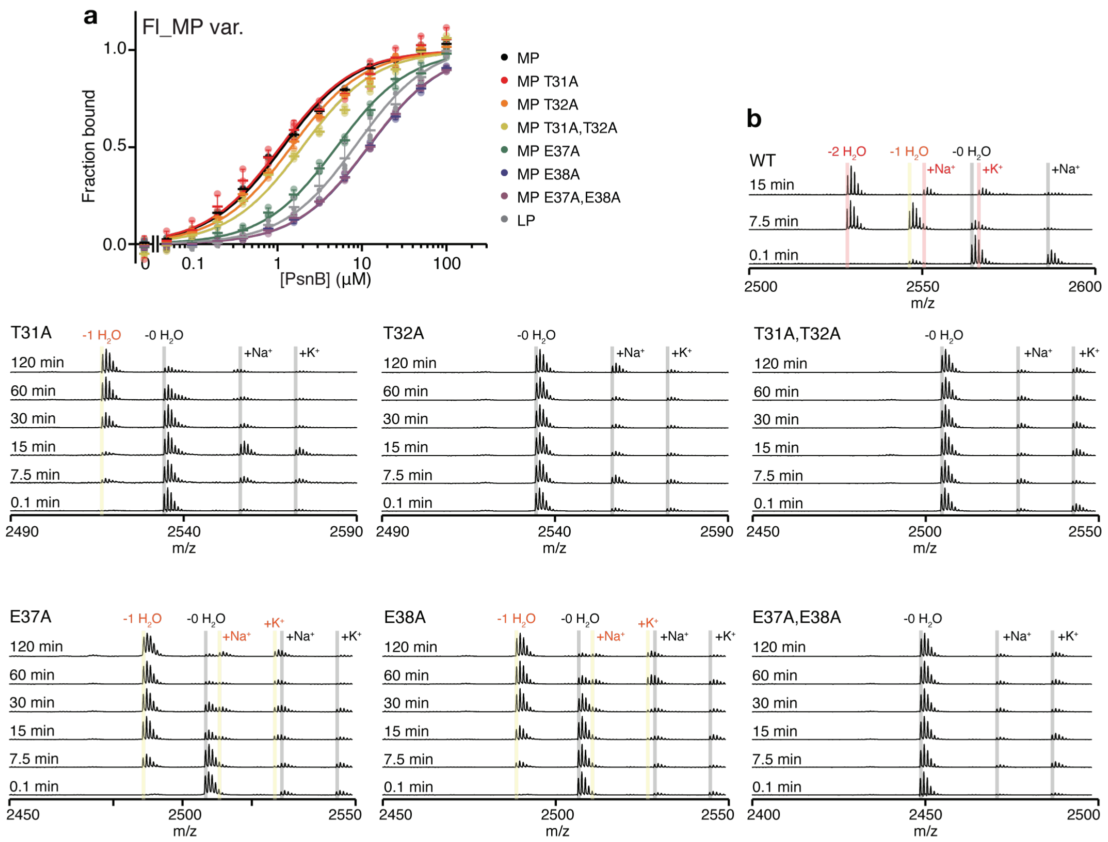


**Supplementary figure 4. Binding and modification property of MP variants. a**, Affinity of Fl_MP variants (0.1 μM) to PsnB was measured by fluorescence anisotropy. Data are presented as dot plots with mean ±1 SD (n = 3 independent experiments) and fitted to a hyperbolic equation. **b**, Macrolactone formation of MP variants by the PsnB was monitored by MALDI. 1 μM PsnB and 50 μM MP variants were incubated in buffer A at 37°C, and the reaction solutions at designated time points were analyzed by MALDI-MS.

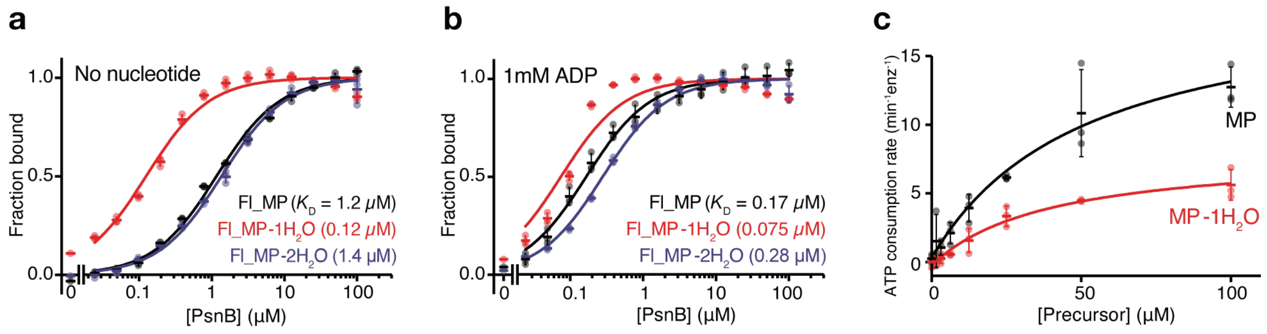


**Supplementary figure 5. Binding and modification of ring-containing precursors.** **a**,**b**, Affinity of ring-containing MPs (MP-1$H_2O$, the single-ring intermediate; MP-2$H_2O$, the double-ring product) to PsnB was determined by fluorescence anisotropy without nucleotide (**a**) or with 1 mM ADP (**b**). **c**, ATPase activity of PsnB was measured with different concentrations of MP or the single-ring intermediate (MP-1$H_2O$). Data are presented as dot plots with mean ±1 SD (n = 3 independent experiments; **a-c**) and fitted to a hyperbolic equation (**a-c**).

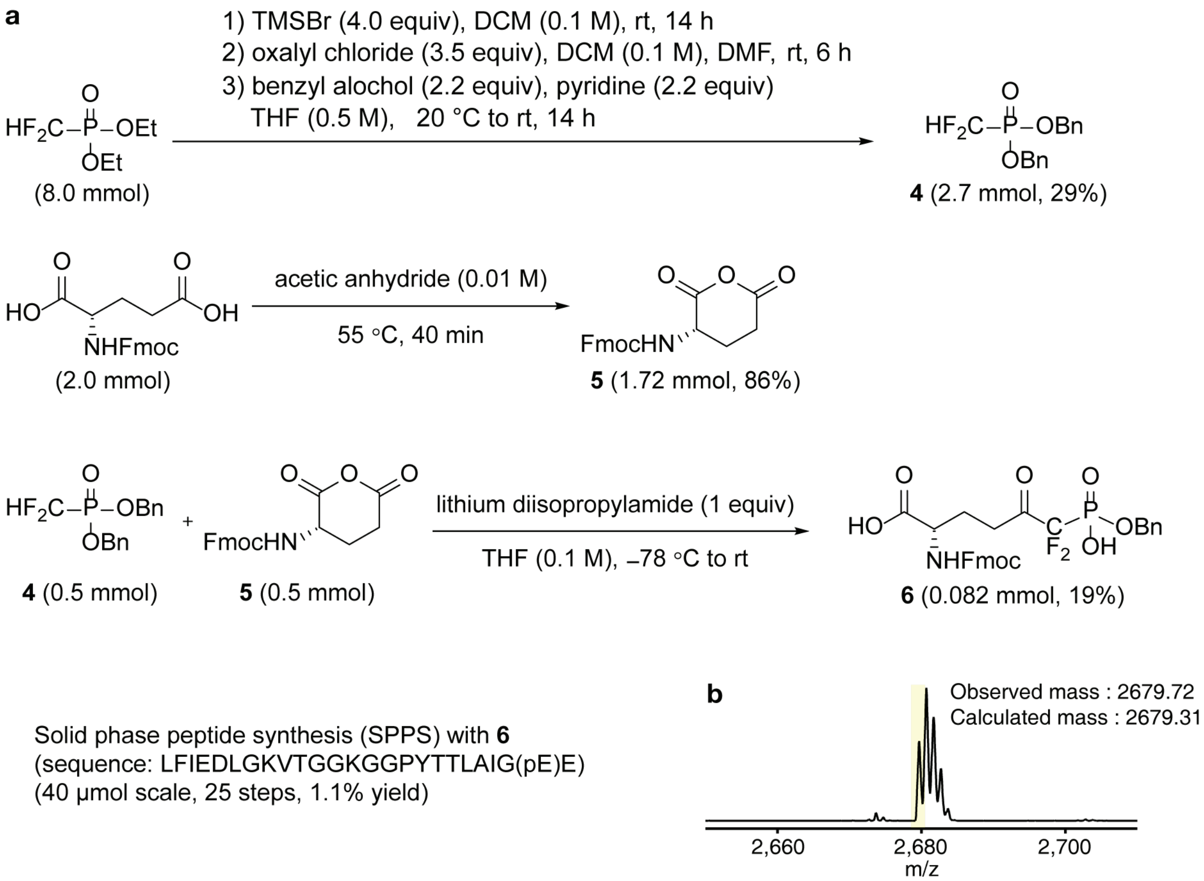


**Supplementary figure 6. Chemical synthesis of a phosphomimetic glutamate (6). a,** Phosphomimetic glutamate (pE) is synthesized as following scheme. Dibenzyl-difluoromethylphosphonate **4** was generated from Diethyl-difluoromethylphosphonate. Fmoc-protected glutamic anhydride **5** was generated from Fmoc-protected glutamic acid[2]. From **4** and **5**, Fmoc-protected benzyl-difluoromethylphosphonate **6** was furnished by using LDA[3]. Detailed synthesis procedure is described in Supplementary Note. With **6**, peptide containing pE was synthesized with the yield of 1.1%. Peptide synthesis was performed as described in the method. **b**, MALDI spectrum of synthesized phosphomimetic variant at Glu37(MP(pE37)).

**a**

| PDB code (resolution) | Complex | State | Assembly | PsnB chain | Precursor chain | Nucleotide | Cα RMSD (# of Cα) | Loop β6α3 | Loop β13β14 | Resolved region of precursor |
|---|---|---|---|---|---|---|---|---|---|---|
| 7DRM (2.75 Å) | PsnB PsnA2$_{14-38}$ ADP | ENLC-E | ABE | A | E | ADP | 1.556 (293) | O | O | 14-22(LP)/33-38(CP) |
| | | | | B | - | - | | X | X | - |
| | | | CDF | C | F | ADP | 1.378 (292) | O | O | 14-22(LP)/32-38(CP) |
| | | | | D | - | - | | X | X | - |
| 7DRN (3.14 Å) | PsnB PsnA2$_{14-38}$ AMPPNP | | ABE | A | E | AMPPNP | 1.458 (292) | O | O | 14-22(LP)/31-38(CP) |
| | | | | B | - | - | | X | X | - |
| | | | CDF | C | F | AMPPNP | 1.395 (292) | O | O | 14-22(LP)/31-38(CP) |
| | | | | D | - | - | | X | X | - |
| 7DRP (2.63 Å) | PsnB PsnA2$_{14-38}$ (pE37) ADP | ENLC-EN | ABE | A | E | ADP | 1.484 (294) | O | O | 14-21(LP)/32-38(CP) |
| | | | | B | - | ADP | | X | X | - |
| | | | CDF | C | F | ADP | 1.338 (291) | O | O | 14-21(LP)/29-38(CP) |
| | | | | D | - | ADP | | X | X | - |
| 7DRO (2.95 Å) | PsnB PsnA2$_{14-38}$ | EL-EL | ABGH | A | G | - | 0.541 (297) | X | O | 14-19(LP) |
| | | | | B | H | - | | X | O | 14-19(LP) |
| | | EL-E | CDI | C | I | - | 0.803 (283) | X | O | 14-19(LP) |
| | | | | D | - | - | | X | X | - |
| | | | EFJ | E | J | - | 0.888 (280) | X | O | 14-19(LP) |
| | | | | F | - | - | | X | X | - |

**b**

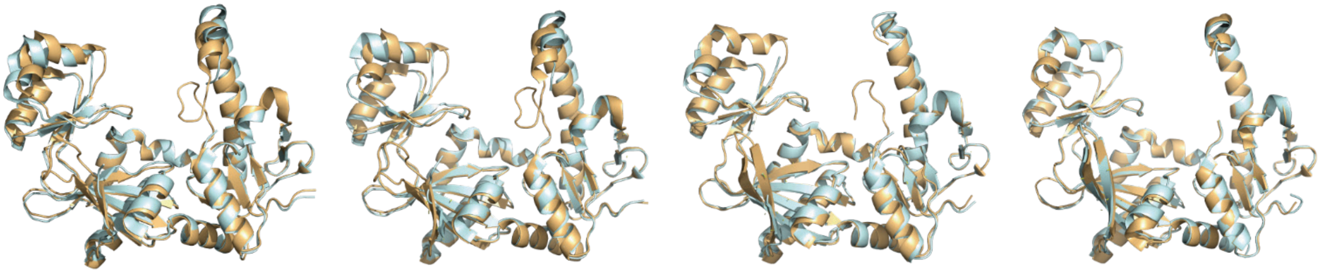

Overlap of Chain A and B in 7DRM RMSD = 1.556 Å for Cα number of 293 (ENLC-E)

Overlap of Chain A and B in 7DRP RMSD = 1.484 Å for Cα number of 294 (ENLC-EN)

Overlap of Chain C and D in 7DRO RMSD = 0.803 Å for Cα number of 283 (EL-E)

Overlap of Chain A and B in 7DRO RMSD = 0.541 Å for Cα number of 297 (EL-EL)

**Supplementary figure 7. Four crystal structures of PsnB complexes.** **a**, Basic information for four crystal structures of PsnB complexes. 7DRM, 7DRN, and 7DRP contain two PsnB dimers in the asymmetric unit, whereas 7DRO has three PsnB dimers. Total nine independent dimers were classified into four states based on the components of the dimer. Four states of PsnB dimers show different level of asymmetry between two monomers and different loop stability (O: resolved, X: not resolved). **b**, Two superposed PsnB subunits in the dimers (yellow and light blue cartoons). Root-mean-square deviations (RMSDs) of Cα atoms are shown below.

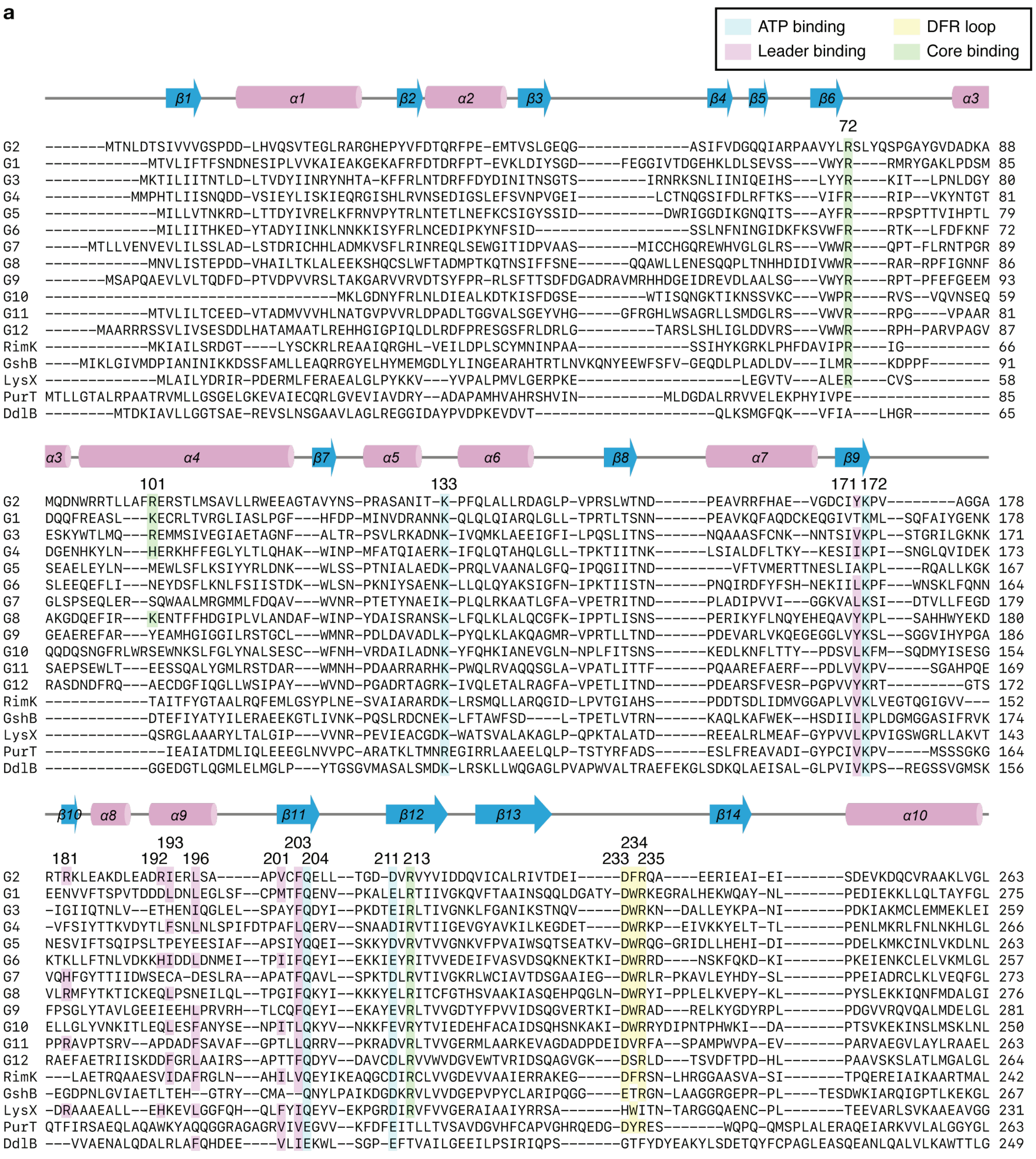
a
ATP binding
Leader binding
DFR loop
Core binding
β1 α1 β2 α2 β3 β4 β5 β6 α3
72
G2 -------MTNLDTSIVVVGSPDD-LHVQSVTEGLRARGHEPYVFDTQRFPE-EMTVSLGEQG-------------ASIFVDGGQQIARPAAVYLRSLYQSPGAYGVDADKA 88
G1 ------------MTVLIFTFSNDNESIPLVVKAIEAKGEKAFRFDTDRFPT-EVKLDIYSGD-----FEGGIVTDGEHKLDLSEVSS---VWYR----RMRYGAKLPDSM 85
G3 -----------MKTILIITNTLD-LTVDYIINRYNHTA-KFFRLNTDRFFDYDINITNSGTS--------IRNRKSNLIINIQEIHS---LYYR----KIT--LPNLDGY 80
G4 ----------MMPHTLIISNQDD-VSIEYLISKIEQRGISHLRVNSEDIGSLEFSVNPVGEI---------LCTNQGSIFDLRFTKS---VIFR----RIP--VKYNTGT 81
G5 -------------MILLVTNKRD-LTTDYIVRELKFRNVPYTRLNTETLNEFKCSIGYSSID----------DWRIGGDIKGNQITS---AYFR----RPSPTTVIHPTL 79
G6 -------------MILIITHKED-YTADYIINKLNNKKISYFRLNCEDIPKYNFSID------------------SSLNFNINGIDKFKSVWFR----RTK--LFDFKNF 72
G7 -----MTLLVENVEVLILSSLAD-LSTDRICHHLADMKVSFLRINREQLSEWGITIDPVAAS-------MICCHGQREWHVGLGLRS---VWWR----QPT-FLRNTPGR 89
G8 ------------MNVLISTEPDD-VHAILTKLALEEKSHQCSLWFTADMPTKQTNSIFFSNE------QQAWLLENESQQPLTNHHDIDIVWWR----RAR-RPFIGNNF 86
G9 -------MSAPQAEVLVLTQDFD-PTVDPVVRSLTAKGARVVRVDTSYFPR-RLSFTTSDFDGADRAVMRHHDGEIDREVDLAALSG---VWYR----RPT-PFEFGEEM 93
G10 ----------------------------------MKLGDNYFRLNLDIEALKDTKISFDGSE--------WTISQNGKTIKNSSVKC---VWPR----RVS--VQVNSEQ 59
G11 ------------MTVLILTCEED-VTADMVVVHLNATGVPVVRLDPADLTGGVALSGEYVHG-----GFRGHLWSAGRLLSMDGLRS---VWVR----RPG----VPAAR 81
G12 ------MAARRRSSVLIVSESDDLHATAMAATLREHHGIGPIQLDLRDFPRESGSFRLDRLG---------TARSLSHLIGLDDVRS---VWWR----RPH-PARVPAGV 87
RimK ------------MKIAILSRDGT----LYSCKRLREAAIQRGHL-VEILDPLSCYMNINPAA-------------SSIHYKGRKLPHFDAVIPR----IG---------- 66
GshB ----MIKLGIVMDPIANINIKKDSSFAMLLEAQRRGYELHYMEMGDLYLINGEARAHTRTLNVKQNYEEWFSFV-GEQDLPLADLDV---ILMR----KDPPF------- 91
LysX -------------MLAILYDRIR-PDERMLFERAEALGLPYKKV---YVPALPMVLGERPKE------------------LEGVTV---ALER----CVS--------- 58
PurT MTLLGTALRPAATRVMLLGSGELGKEVAIECQRLGVEVIAVDRY--ADAPAMHVAHRSHVIN----------MLDGDALRRVVELEKPHYIVPE---------------- 85
DdlB --------MTDKIAVLLGGTSAE-REVSLNSGAAVLAGLREGGIDAYPVDPKEVDVT--------------------QLKSMGFQK---VFIA---LHGR--------- 65
α3 α4 β7 α5 α6 β8 α7 β9
101 133 171 172
G2 MQDNWRRTLLAFRERSTLMSAVLLRWEEAGTAVYNS-PRASANIT-K-PFQLALLRDAGLP-VPRSLWTND------PEAVRRFHAE--VGDCIYKPV--------AGGA 178
G1 DQQFREASL---KECRLTVRGLIASLPGF---HFDP-MINVDRANNK-QLQLQIARQLGLL-TPRTLTSNN------PEAVKQFAQDCKEQGIVTKML--SQFAIYGENK 178
G3 ESKYWTLMQ---REMMSIVEGIAETAGNF---ALTR-PSVLRKADNK-IVQMKLAEEIGFI-LPQSLITNS------NQAAASFCNK--NNTSIVKPL--STGRILGKNK 171
G4 DGENHKYLN---HERKHFFEGLYLTLQHAK--WINP-MFATQIAERK-IFQLQTAHQLGLL-TPKTIITNK------LSIALDFLTKY--KESIIKPI--SNGLQVIDEK 173
G5 SEAELEYLN---MEWLSFLKSIYYRLDNK---WLSS-PTNIALAEDK-PRQLVAANALGFQ-IPQGIITND---------VFTVMERTTNESLIAKPL----RQALLKGK 167
G6 SLEEQEFLI---NEYDSFLKNLFSIISTDK--WLSN-PKNIYSAENK-LLQLQYAKSIGFN-IPKTIISTN------PNQIRDFYFSH-NEKIILKPF---WNSKLFQNN 164
G7 GLSPSEQLER--SQWAALMRGMMLFDQAV---WVNR-PTETYNAEIK-PLQLRKAATLGFA-VPETRITND------PLADIPVVI---GGKVALKSI---DTVLLFEGD 179
G8 AKGDQEFIR---KENTFFHDGIPLVLANDAF-WINP-YDAISRANSK-LFQLKLALQCGFK-IPPTLISNS------PERIKYFLNQYEHEQAVYKPL---SAHHWYEKD 180
G9 GEAEREFAR---YEAMHGIGGILRSTGCL---WMNR-PDLDAVADLK-PYQLKLAKQAGMR-VPRTLLTND------PDEVARLVKQEGEGGLVYKSL--SGGVIHYPGA 186
G10 QQDQSNGFRLWRSEWNKSLFGLYNALSESC--WFNH-VRDAILADNK-YFQHKIANEVGLN-NPLFITSNS------KEDLKNFLTTY--PDSVLKFM--SQDMYISESG 154
G11 SAEPSEWLT---EESSQALYGMLRSTDAR---WMNH-PDAARRARHK-PWQLRVAQQSGLA-VPATLITTF------PQAAREFAERF--PDLVVKPV-----SGAHPQE 169
G12 RASDNDFRQ---AECDGFIQGLLWSIPAY---WVND-PGADRTAGRK-IVQLETALRAGFA-VPETLITND------PDEARSFVESR-PGPVVYKRT---------GTS 172
RimK ------------TAITFYGTAALRQFEMLGSYPLNE-SVAIARARDK-LRSMQLLARQGID-LPVTGIAHS------PDDTSDLIDMVGGAPLVVKLVEGTQGIGVV--- 152
GshB ------------DTEFIYATYILERAEEKGTLIVNK-PQSLRDCNEK-LFTAWFSD----L-TPETLVTRN------KAQLKAFWEK--HSDIILKPLDGMGGASIFRVK 174
LysX ------------QSRGLAAARYLTALGIP---VVNR-PEVIEACGDK-WATSVALAKAGLP-QPKTALATD------REEALRLMEAF-GYPVVLKPVIGSWGRLLAKVT 143
PurT --------------IEAIATDMLIQLEEEGLNVVPC-ARATKLTMNREGIRRLAAEELQLP-TSTYRFADS------ESLFREAVADI-GYPCIVKPV-----MSSSGKG 164
DdlB ------------GGEDGTLQGMLELMGLP---YTGSGVMASALSMDK-LRSKLLWQGAGLPVAPWVALTRAEFEKGLSDKQLAEISAL-GLPVIVKPS--REGSSVGMSK 156
β10 α8 α9 β11 β12 β13 β14 α10
193 203 234
181 192 196 201 204 211 213 233 235
G2 RTRKLEAKDLEADRIERLSA-----APVCFQELL--TGD-DVRVYVIDDQVICALRIVTDEI-----DFRQA----EERIEAI-EI-------SDEVKDQCVRAAKLVGL 263
G1 EENVVFTSPVTDDDLDNLEGLSF--CPMTFQENV--PKALELRTIIVGKQVFTAAINSQQLDGATY-DWRKEGRALHEKWQAY-NL-------PEDIEKKLLQLTAYFGL 275
G3 -IGIIQTNLV--ETHENIQGLEL--SPAYFQDYI--PKDTEIRLTIVGNKLFGANIKSTNQV-----DWRKN--DALLEYKPA-NI-------PDKIAKMCLEMMEKLEI 259
G4 -VFSIYTTKVDYTLFSNLNLSPIFDTPAFLQERV--SNAADIRVTIIGEVGYAVKILKEGDET----DWRKP--EIVKKYELT-TL-------PENLMKRLFNLNKHLGL 266
G5 NESVIFTSQIPSLTPEYEESIAF--APSIYQQEI--SKKYDVRVTVVGNKVFPVAIWSQTSEATKV-DWRQG-GRIDLLHEHI-DI-------PDELKMKCINLVKDLNL 263
G6 KTKLLFTNLVDKKHIDDLDNMEI--TPIIFQEYI--EKKIEYRITVVEDEIFVASVDSQKNEKTKI-DWRRD----NSKFQKD-KI-------PKEIENKCLELVKMLGL 257
G7 VQHFGYTTIIDWSECA-DESLRA--APATFQAVL--SPKTDLRVTIVGKRLWCIAVTDSGAAIEG--DWRLR-PKAVLEYHDY-SL-------PPEIADRCLKLVEQFGL 273
G8 VLRMFYTKTICKEQLPSNEILQL--TPGIFQKYI--KKKYELRITCFGTHSVAAKIASQEHPQGLN-DWRYI-PPLELKVEPY-KL-------PYSLEKKIQNFMDALGI 276
G9 FPSGLYTAVLGEEIEEHLPRVRH--TLCQFQEYI--EKAYEVRLTVVGDTYFPVVIDSQGVERTKI-DWRAD---RELKYGDYRPL-------PDGVVRQVQALMDELGL 281
G10 ELLGLYVNKITLEQLESFANYSE--NPITLQKYV--NKKFEVRYTVIEDEHFACAIDSQHSNKAKI-DWRRYDIPNTPHWKI--DA-------PTSVKEKINSLMSKLNL 250
G11 PPRAVPTSRV--APDADFSAVAF--GPTLLQRRV--PKRADVRLTVVGERMLAARKEVAGDADPDEIDVRFA--SPAMPWVPA-EV-------PARVAEGVLAYLRAAEL 263
G12 RAEFAETRIISKDDFGRLAAIRS--APTTFQDYV--DAVCDLRVVWVDGVEWTVRIDSQAGVGK---DSRLD---TSVDFTPD-HL-------PAAVSKSLATLMGALGL 264
RimK ---LAETRQAAESVIDAFRGLN---AHILVQEYIKEAQGCDIRCLVVGDEVVAAIERRAKEG-----DFRSN-LHRGGAASVA-SI-------TPQEREIAIKAARTMAL 242
GshB -EGDPNLGVIAETLTEH--GTRY--CMA--QNYLPAIKDGDKRVLVVDGEPVPYCLARIPQGG----ETRGN-LAAGGRGEPR-PL----TESDWKIARQIGPTLKEKGL 267
LysX -DRAAAEALL--EHKEVLGGFQH--QLFYIQEYV-EKPGRDIRVFVVGERAIAAIYRRSA-------HWITN-TARGGQAENC-PL-------TEEVARLSVKAAEAVGG 231
PurT QTFIRSAEQLAQAWKYAQQGGRAGAGRVIVEGVV--KFDFEITLLTVSAVDGVHFCAPVGHRQEDG-DYRES-------WQPQ-QMSPLALERAQEIARKVVLALGGYGL 263
DdlB ---VVAENALQDALRLAFQHDEE----VLIEKWL--SGP-EFTVAILGEEILPSIRIQPS-------GTFYDYEAKYLSDETQYFCPAGLEASQEANLQALVLKAWTTLG 249

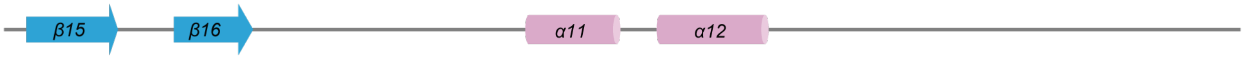


```
          269            282 284
G2   R--YTGMDIKAGADGNYRVLELNAS--------------AM------FRGFEGRANV-DICGPLCDALIAQTKR------------------------------------- 314
G1   N--YGAIDIIVTPDGRHVFLEINPV--------------GE------FFWLELFAPYFPISQAIAEILLTPTNQKPF---------------------------------- 330
G3   N--FAAFDFII-RNGDYIFLELNAN--------------GQ------WLWLEDILKF-DISNTIINYLLGEPI---------------------------------------- 308
G4   V--YSAIDLILTPSGDYVFLEVNPV--------------GE------WVWLELELGI-CISEQILKELL-------------------------------------------- 312
G5   R--FGAIDLVCDLDGDYWFLEINPN--------------GQ------WAWIENQTKL-PIASSIVDELLKISENNYES----------------------------------- 318
G6   K--FGAIDLIQDEYDNFYFLEINAN--------------GQ------WVWIESDTGL-TISDSIINFLNA------------------------------------------- 304
G7   E--YAAIDLALSND-QYWFIEINPT--------------GE------WGWLDRGGR--GISNAIAEALACRGQPS-------------------------------------- 323
G8   V--FGCFDFIVTPEGDYYFLEVNQQ--------------GQ------FLWIEDCCPTIKMLDKFVKFILNRDAYFNWKDKETISLADYEEKAAVVVYENMQKHIMLNALK 364
G9   V--YAALDFIVTPDGEHVFLESNPN--------------GQ------FMWMQHDLGL-PMADRIADLLMRGGPFRRGDVEQVGY-------------------------- 342
G10  R--YGALDFIVDENNDWWFLEVNSA--------------GQ------WLWIEDLSGM-DISGSIANWLTSNSR--------------------------------------- 300
G11  T--YGAFDFAEDSDGTWWFLECNQS--------------GQ------FGFVEVETGQ-PIARTIAEWLAAPGAEERESCDGSDSVVP----------------------- 327
G12  S--FGVLDIRLGRDGEYYFLEVNPQ--------------GQ------FAYLEVKTGL-PIFRSLANLLVDGDGTVAGY---------------------------------- 319
RimK D--VAGVDILRANRGP-LVMEVNAS--------------PG------LEGIEKTTGI-DIAGKMIRWIERHATTEYCLKTGG------------------------------ 300
GshB I--FVGLDIIGDR-----LTEINVTSP------------TC------IREIEAEFPV-SITGMLMDAIEARLQQQ-------------------------------------- 316
LysX G--VVAVDLFESERG-LLVNEVNHT--------------ME------FKNSVHTTGV-DIPGEILKYAWSLAS---------------------------------------- 280
PurT ---FGV-ELFVC-GDEVIFSEVSPRPHDTGMVTLISQDLSE-----FALHVRAFLGL-PVGGIRQYGPAASAVILPQLTSQNVTFDNVQNAVGADLQIRLFGKPEIDGSR 362
DdlB CKGWGRIDVMLDSDGQFYLLEANTS--------------PGMTSHSLVPMAARQAGM-SFSQLVVRILELAD----------------------------------------- 306
```

```
G2   ------------------------------ 314
G1   ------------------------------ 330
G3   ------------------------------ 308
G4   ------------------------------ 312
G5   ------------------------------ 318
G6   ------------------------------ 304
G7   ------------------------------ 323
G8   KTEAA------------------------- 369
G9   ------------------------------ 342
G10  ------------------------------ 300
G11  ------------------------------ 327
G12  ------------------------------ 319
RimK ------------------------------ 300
GshB ------------------------------ 316
LysX ------------------------------ 280
PurT RLGVALATAESVVDAIERAKHAAGQVKVQG 392
DdlB ------------------------------ 306
```

**b**

| | Graspetide | | | | | | | | | | | | | | | | | | | | |
|---|---|---|---|---|---|---|---|---|---|---|---|---|---|---|---|---|---|---|---|---|---|
| | G2 (PsnB) | Residue number | Ref | G2a (42) | G2 (202) | G1 | G3 | G4 | G5 | G6 | G7 | G8 | G9 | G10 | G11 | G12 | Grasptide (2024) | RimK | GshB | LysX | PurT | DdlB |
| ATP binding | K | 133 | K | 100 | 100 | K | K | K | K | K | K | K | K | K | K | K | 99.4 | K | K | K | R | K |
| | K | 172 | K | 100 | 100 | K | K | K | K | K | K | K | K | K | K | K | 99.9 | K | K | K | K | K |
| | Q | 204 | Q | 100 | 100 | Q | Q | Q | Q | Q | Q | Q | Q | Q | Q | Q | 99.8 | Q | Q | Q | E | E |
| | D | 211 | D/E | 97.6 | 99.5 | E | E | D | D | E | D | E | E | E | D | D | 99.6 | D | D | D | E | E |
| | D | 269 | D | 100 | 99.5 | D | D | D | D | D | D | D | D | D | D | D | 99.2 | D | D | D | E | D |
| | E | 282 | E | 97.6 | 99 | E | E | E | E | E | E | E | E | E | E | E | 98.8 | E | E | E | E | E |
| | N | 284 | N | 100 | 99.5 | N | N | N | N | N | N | N | N | N | N | N | 98.9 | N | N | N | S | N |
| DFR | D | 233 | D/N | 100 | 100 | D | D | D | D | D | D | D | D | D | D | D | 99.7 | D | E | H | D | G |
| | F | 234 | aromatic | 97.6 | 99 | W | W | W | W | W | W | W | W | W | V | S | 71.1 | F | T | W | Y | T |
| | R | 235 | R | 100 | 100 | R | R | R | R | R | R | R | R | R | R | R | 99.6 | R | R | I | R | F |
| Leader binding | Y | 171 | Y | 90.4 | 43.6 | T | V | I | A | L | L | Y | Y | L | V | Y | 12.1 | V | L | L | V | V |
| | I | 193 | I/L | 85.7 | 90.1 | L | I | L | E | L | D | N | L | F | F | L | 49.2 | F | - | L | Q | F |
| | L | 196 | L | 92.9 | 72.8 | L | L | I | I | M | L | L | V | Y | V | I | 42.3 | L | H | F | A | Q |
| | V | 201 | V | 95.2 | 69.8 | M | A | A | S | I | A | G | C | I | T | T | 12.2 | I | A | F | V | V |
| | F | 203 | F | 95.2 | 49.5 | F | F | L | Y | F | F | F | F | L | L | F | 52 | V | A | I | V | I |
| | R | 181 | R | 64.3 | 25.2 | N | G | F | S | K | H | R | S | L | R | E | 20.2 | - | G | R | F | - |
| | R | 192 | R | 90.5 | 44.6 | D | T | L | T | H | E | Q | I | Q | P | D | 5.5 | V | L | H | W | L |
| Core binding | R | 72 | R | 100 | 71.8 | R | R | R | R | R | R | R | R | R | R | R | 96.3 | R | R | R | E | A |
| | R | 213 | R | 100 | 100 | R | R | R | R | R | R | R | R | R | R | R | 99.8 | R | R | R | T | T |
| | R | 101 | R | 97.6 | 30.7 | K | R | H | M | N | S | K | Y | S | E | A | 14 | T | D | Q | - | G |

**Supplementary figure 8. Conservation pattern of residues implicated in substrate interaction of ATP-grasp enzymes. a**, Sequence alignment of PsnB (an enzyme for Group 2 graspetide, G2), representative enzymes for remaining 11 graspetide groups (G1 and G3-G12), and non-RiPP ATP-grasp enzymes (RimK, GshB, LysX, PurT, and DdlB). Secondary structures of PsnB are shown above the alignment. Critical residues for specific interaction are highlighted (ATP binding, cyan; DFR loop, yellow; leader binding, magenta; core binding, green). Strain names and NCBI accession numbers of the representative enzymes for 12 graspetides and non-RiPP ATP-grasp enzymes are as follows: G1, *Planktothrix agardhii*, WP_042156020.1; G2, *Plesiocystis pacifica*, WP_006971586.1; G3, *Bacillus thuringiensis*, WP_000849148.1; G4, *Sphingobacteriales bacterium* 44-61, OJW02008.1; G5, *Vibrio* sp. JCM 18905, GAJ78971.1; G6, *Chryseobacterium greenlandense*, WP_059136627.1; G7, *Agrobacterium* sp. SUL3, WP_052821370.1; G8, *Legionella beliardensis*, WP_115303353.1; G9, *Actinomadura darangshiensis*, WP_132205758.1; G10, *Citrobacter* sp. wls827, WP_137346417.1; G11, *Streptomyces acidiscabies*, WP_059044944.1; G12, *Nocardia abscessus*, WP_043693582.1; RimK, *Escherichia coli* K-12, P0C0U4.1; GshB, *Escherichia coli* K-12, P04425.1; LysX, *Thermus thermophilus* HB8, Q5SH23; PurT, *Escherichia coli* K-12, P33221; DdlB, *Escherichia coli* K-12, P07862. **b**, Summary of the conservation pattern of critical residues. Yellow and light yellow indicate identical and synonymous residues, respectively. Conservation % in Group 2a or 2 graspetides or in all graspetides are also shown with the number of members in each group (70-90% conservation, light green; 90-100%, green).

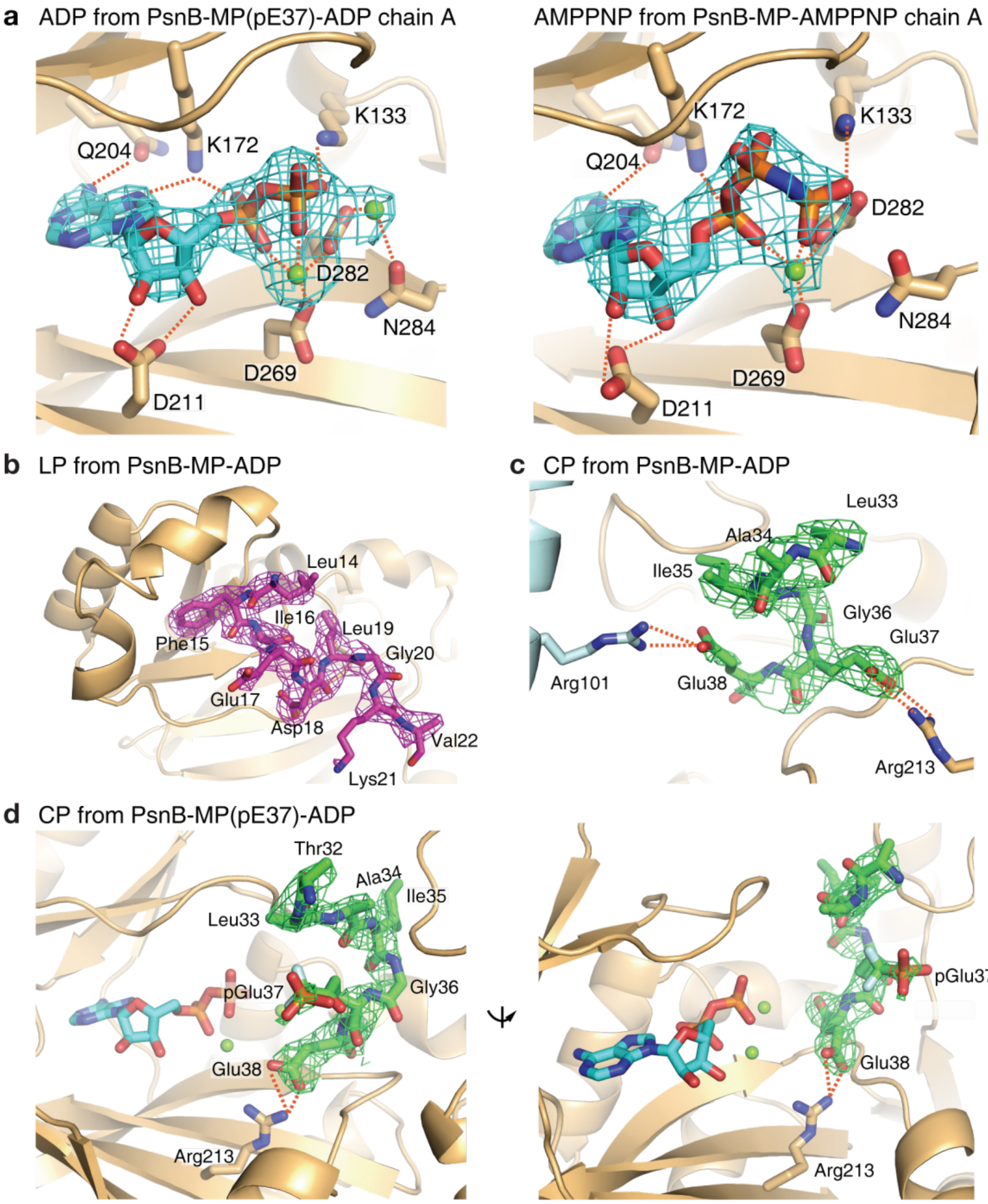


**Supplementary figure 9. Electron density maps for nucleotide, LP, and CP. a**, Modeled ADP and AMPPNP (sticks) are shown with the calculated 2mF$_o$-DF$_c$ electron density map displayed at 2.0σ and 1.5σ contour level, respectively. PsnB subunit (yellow cartoon) and the highly conserved residues at the ATP-binding site (yellow sticks) are also shown. **b**, LP (magenta sticks) in the PsnB-MP-ADP complex (7DRM) is shown with the calculated 2mF$_o$-DF$_c$ electron density map displayed at 1.0σ contour level. Highly conserved LFIEDL residues and additional GKV residues of LP are well resolved in the electron density map. **c**, CP (green sticks) in the PsnB-MP-ADP complex (7DRM) is shown with the calculated 2mF$_o$-DF$_c$ electron density map displayed at 1.0σ contour level. Two arginines (Arg213 and Arg101; yellow or light blue sticks) that interact with two glutamates in CP are also shown. Majority of core peptides (LAIGEE) are well resolved in the electron density map. **d**, CP (green sticks) in the PsnB-MP(pE37)-ADP complex (7DRP) is shown with the calculated 2mF$_o$-DF$_c$ electron density map displayed at a 1.0σ contour level. Majority of core residues (TLAIG(pE)E) containing phosphomimetic residue (pGlu37) are well resolved in the electron density map. pGlu37 makes no interaction, whereas Glu38 interacts with Arg213 (yellow sticks) of PsnB.

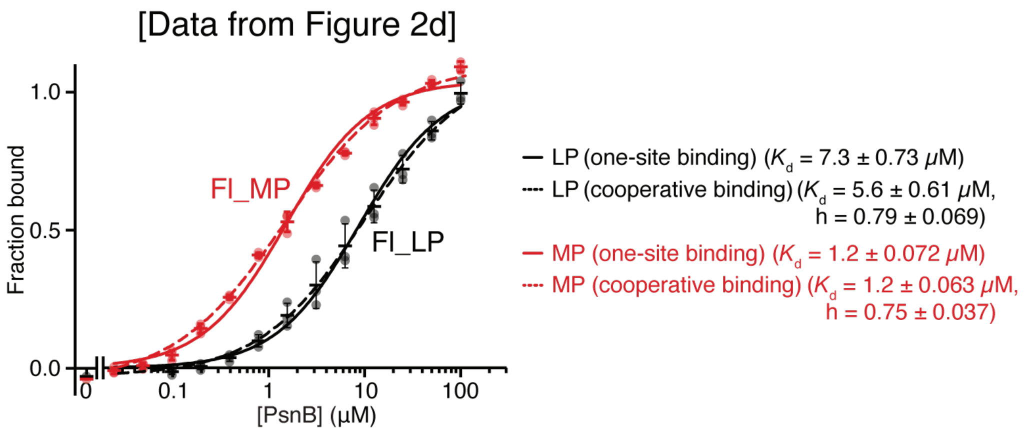


**Supplementary figure 10. Binding of MP and LP to PsnB shows negative cooperativity.** Data in Figure 2d are fitted to two different equations: a normal hyperbolic equation (solid lines) and a Hill equation (dashed lines). The latter fitting resulted in the Hill coefficients of 0.75 and 0.79 for MP and LP, respectively, suggesting the negative cooperativity of the interactions.

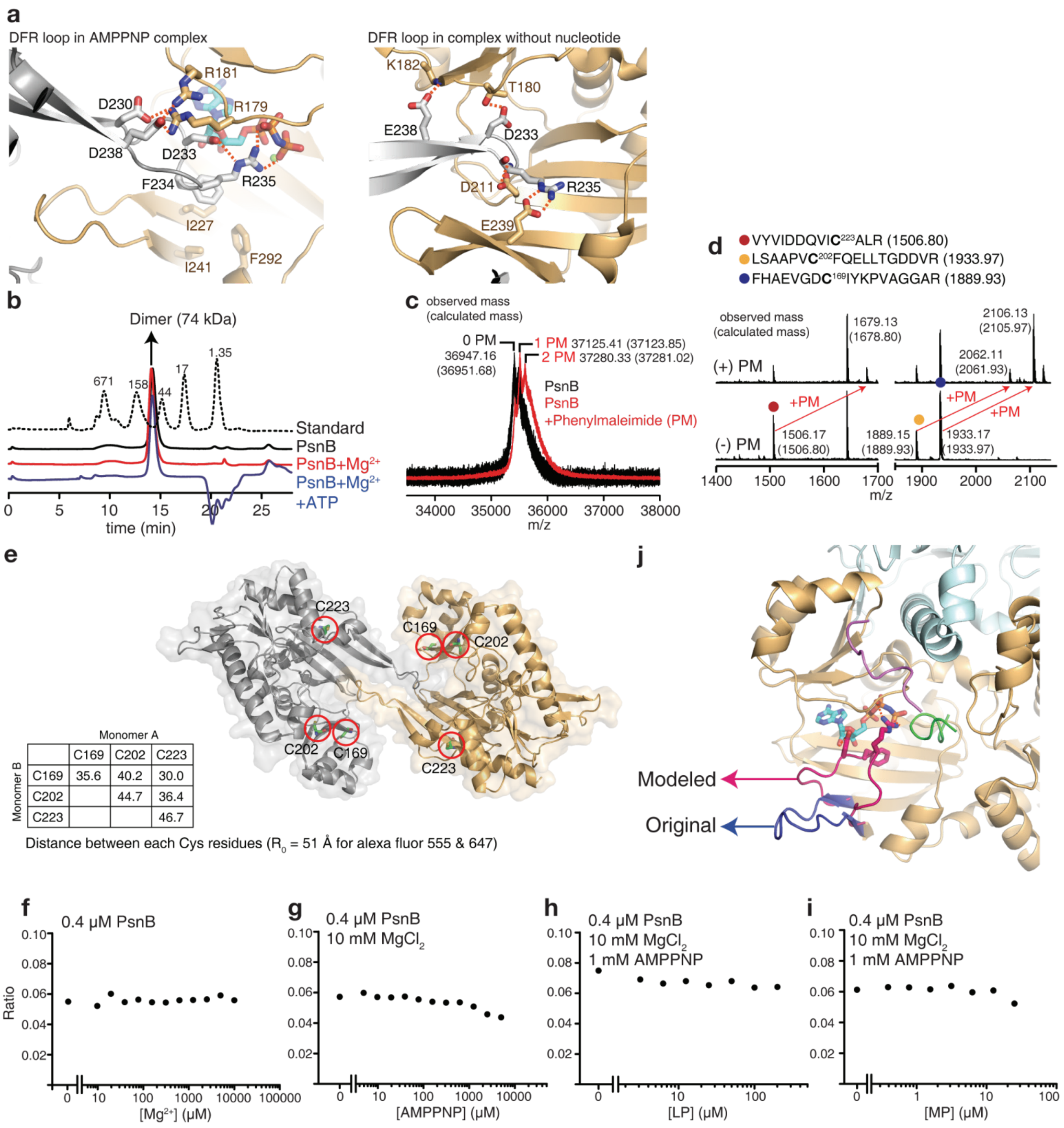
a
DFR loop in AMPPNP complex
R181
D230
R179
D238
D233
F234
R235
I227
F292
I241
DFR loop in complex without nucleotide
K182
T180
E238
D233
D211
R235
E239
b
Dimer (74 kDa)
671
158
44
17
1.35
Standard
PsnB
PsnB+Mg2+
PsnB+Mg2+
+ATP
time (min)
c
observed mass
(calculated mass)
0 PM
36947.16
(36951.68)
1 PM 37125.41 (37123.85)
2 PM 37280.33 (37281.02)
PsnB
PsnB
+Phenylmaleimide (PM)
m/z
d
VYVIDDQVIC223ALR (1506.80)
LSAAPVC202FQELLTGDDVR (1933.97)
FHAEVGDC169IYKPVAGGAR (1889.93)
observed mass
(calculated mass)
(+) PM
1679.13
(1678.80)
2106.13
(2105.97)
2062.11
(2061.93)
+PM
(-) PM
1506.17
(1506.80)
1889.15
(1889.93)
1933.17
(1933.97)
m/z
e
C223
C169
C202
C202
C169
C223
Monomer A
Monomer B
| | C169 | C202 | C223 |
| C169 | 35.6 | 40.2 | 30.0 |
| C202 | | 44.7 | 36.4 |
| C223 | | | 46.7 |
Distance between each Cys residues (R0 = 51 Å for alexa fluor 555 & 647)
j
Modeled
Original
f
0.4 μM PsnB
Ratio
[Mg2+] (μM)
g
0.4 μM PsnB
10 mM MgCl2
[AMPPNP] (μM)
h
0.4 μM PsnB
10 mM MgCl2
1 mM AMPPNP
[LP] (μM)
i
0.4 μM PsnB
10 mM MgCl2
1 mM AMPPNP
[MP] (μM)

**Supplementary figure 11. DFR residues are critical for enzyme activity. a**, Interaction scheme of ENLC-E (left) or EL-EL (right; yellow cartoon/sticks) with DFR loop from a subunit in the neighboring dimer (gray cartoon/sticks). **b**, Gel-filtration chromatogram of PsnB without $Mg^{2+}$ (black solid line), with $Mg^{2+}$ (red solid line), or with both $Mg^{2+}$ and ATP (blue solid line). A chromatogram of molecular weight standard (black dashed line) is also shown with known molecular weights. Addition of nucleotide did not induce the formation of the stable higher oligomer of PsnB dimer. **c**, MALDI-MS spectrum of PsnB or PsnB labeled with phenylmaleimide (PM). 1-2 PM molecules were added to PsnB. **d**, PM-labeled PsnB was cleaved with trypsin and analyzed by MALDI to discover which cysteine residues of PsnB are labeled. Putative labeling sites with neighboring sequences are shown above. **e**, PM was mainly labeled to Cys169, Cys202, and Cys223, which show closer distances between two neighboring dimers (gray and yellow cartoons) than $R_0$ for Alexa fluor 555 and Alexa fluor 647 (51 Å). Pair-wise distances of three Cys residues between two neighboring dimers are shown in a table. PsnB was labeled with fluorescent dye maleimide (Alexa fluor 555 or Alexa fluor 647) with the same condition. After purification with gel-filtration, dye-labeled PsnB was used for FRET experiments. **f**–**i**, FRET of a solution containing both donor- and acceptor-labeled PsnB was measured with the titration of $Mg^{2+}$ (**f**), AMPPNP (**g**), LP (**h**), and MP (**i**). Additional components in solutions are listed above the plots. Neither nucleotide nor precursor induced stable intermolecular interaction of PsnB. (**j**) DFR loop is long and flexible enough for intramolecular interaction. DFR loop region, from Ile227 to Ile247, was modeled by FALC[4, 5] and overall complex structure was optimized by relaxation. In the modeling structure, the conformation of modeled loop was flipped and DFR residues moved toward the enzyme active site. Also, Arg235 had hydrogen bonding with nucleotide which is similar to the intermolecular interaction scheme shown in crystal structures (**a**).

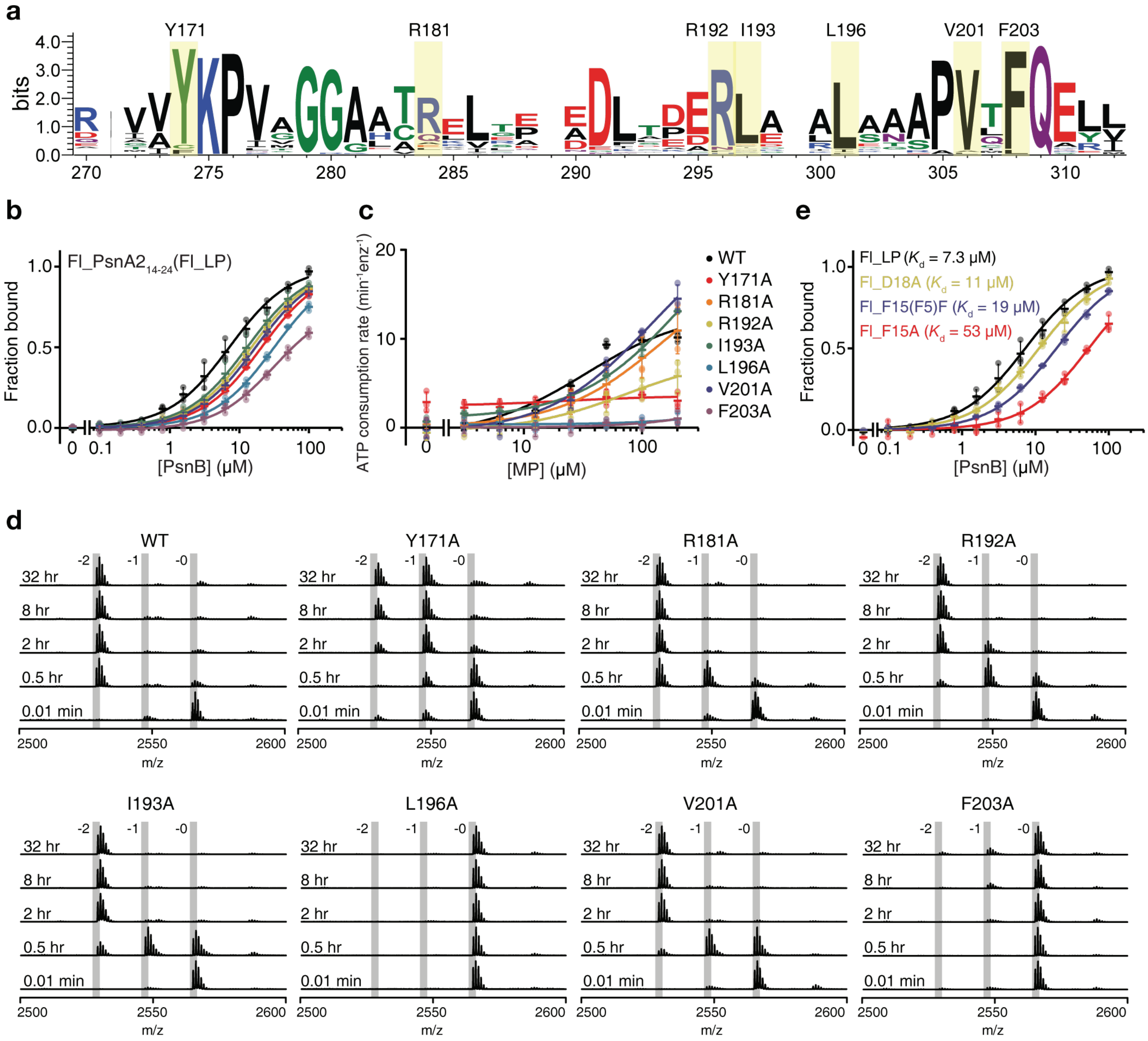


**Supplementary figure 12. Characterization of leader-binding site mutants of PsnB.** **a**, Sequence logo of the leader binding domain in Group 2a graspetide biosynthetic enzymes. Most residues interacting with leader peptide are highly conserved (yellow boxes). **b**,**c**, Fluorescence anisotropy of Fl_LP (0.1 μM; **b**) and ATP-consumption rate (**c**) were measured with PsnB mutants. L196A and F203A were most deleterious. **d**, MALDI-spectra of reactions of the leader-binding site mutants (0.5 μM) with MP (50 μM). **e**, Fluorescence anisotropy of LP variants (0.1 μM) was measured to determine their PsnB affinity. Data are presented as dot plots with mean ±1 SD (n = 3 independent experiments) and fitted to a hyperbolic equation (**b,c,e**).

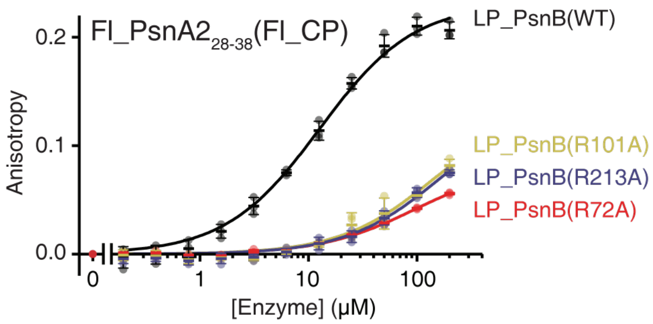


**Supplementary figure 13. Binding property of core-binding site mutants of leader-fused PsnB.** Fluorescence anisotropy of Fl_CP (0.1 μM) with leader-fused PsnB mutants. Mutation of core-binding residues reduced the affinity between CP and the enzyme. Data are presented as dot plots with mean ±1 SD (n = 3 independent experiments) and fitted to a hyperbolic equation.

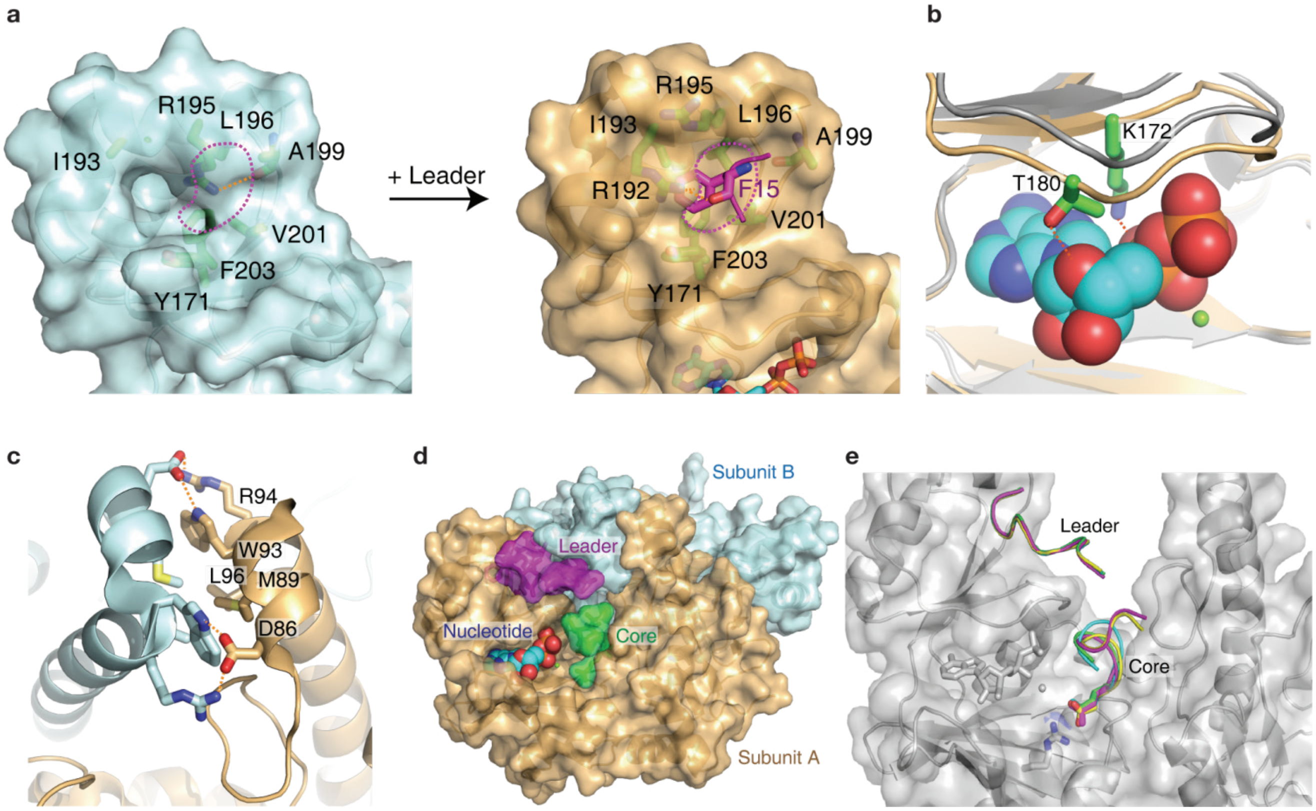


**Supplementary figure 14. Precursor and nucleotide binding induces conformational change of PsnB. a**, Surface models of the leader-binding domain without (left) or with (right) the bound Phe15 in the LP (magenta sticks). Hydrophobic pocket for Phe15 is shown as magenta dashed lines. **b**, The nucleotide (spheres) binding shifts the β9β10 loop (from gray to yellow cartoon). Lys172 and Thr180 that interact with a nucleotide are shown as green sticks. **c**, The two α3α4 pairs of a PsnB dimer show extensive interactions to each other to form a rigid body. Two PsnB subunits are shown as yellow and light blue cartoons, and their interacting residues are presented as yellow and light blue sticks, respectively. **d**, Surface model of the ENLC-E complex (yellow and light blue for two PsnB subunits). Binding of LP (magenta) and nucleotide (spheres) induces the conformational change of PsnB dimer to generate a compact core (green) binding site. **e**, Superposition of four LP and CP pairs from PsnB-MP-ADP (7DRM, green and cyan ribbons) and PsnB-MP-AMPPNP (7DRN, magenta and yellow ribbons). All LPs and AIGEE regions of CP are well overlapped. PsnB is shown as gray cartoon and half-transparent surface. Nucleotide and Arg213 are shown as gray sticks. Glu37 in CP is shown as sticks.

### Supplementary Tables

**Supplementary table 1. Data collection and refinement statistics for 4 structure models**

| | PsnB +PsnA2$_{14-38}$ +ADP | PsnB +PsnA2$_{14-38}$ +AMPPNP | PsnB +PsnA2$_{14-38}$ (pE37) +ADP | PsnB +PsnA2$_{14-38}$ |
|---|---|---|---|---|
| **Data collection** | | | | |
| Space group | P 1 21 1 | P 1 21 1 | P 1 21 1 | P 21 21 21 |
| Cell dimensions | | | | |
| *a*, *b*, *c* (Å) | 91.22, 92.07, 98.83 | 92.29, 92.65, 99.56 | 89.59, 91.66, 98.20 | 86.72, 147.09, 168.18 |
| a, b, g (°) | 90.00, 101.63, 90.00 | 90.00, 100.90, 90.00 | 90.00, 101.31, 90.00 | 90.00, 90.00, 90.00 |
| Resolution (Å) | 33.36-2.75 (2.85-2.75) * | 32.95-3.14 (3.20-3.14) | 33.19-2.63 (2.70-2.63) | 47.08-2.95 (3.00-2.95) |
| $R_{merge}$ | 0.18 (1.92) | 0.25 (1.92) | 0.14 (1.66) | 0.15 (1.46) |
| *I* / s*I* | 5.90 (1.89) | 4.20 (1.79) | 6.80 (2.09) | 6.90 (2.92) |
| Completeness (%) | 99.8 (98.7) | 99.1 (93.3) | 98.9 (91.5) | 99.7 (97.3) |
| Redundancy | 7.30 (7.20) | 7.40 (7.60) | 7.20 (7.30) | 14.3 (11.5) |
| **Refinement** | | | | |
| Resolution (Å) | 33.36-2.75 | 32.95-3.14 | 33.19-2.63 | 47.08-2.95 |
| No. reflections | 41699 | 28824 | 46001 | 45910 |
| $R_{work}$ / $R_{free}$ | 0.226 / 0.242 | 0.248 / 0.263 | 0.216 / 0.238 | 0.249 / 0.293 |
| No. atoms | 9535 | 9516 | 9552 | 13300 |
| Protein | 9479 | 9452 | 9416 | 13300 |
| Ligand/ion | 56 | 90 | 136 | - |
| Water | - | - | - | - |
| *B*-factors | 54.06 | 68.05 | 70.17 | 80.92 |
| Protein | 54.08 | 68.06 | 70.17 | 80.92 |
| Ligand/ion | 50.42 | 66.50 | 69.88 | - |
| Water | - | - | - | - |
| R.m.s. deviations | | | | |
| Bond lengths (Å) | 0.003 | 0.006 | 0.005 | 0.006 |
| Bond angles (°) | 0.77 | 1.15 | 1.12 | 1.12 |

*Single crystal was used for each data set; *Values in parentheses are for highest-resolution shell;

**Supplementary table 2. Plasmid informatics used in this study**

| Plasmids | Description | Reference |
|---|---|---|
| pHB012 | pET28b-6His-[Tm]-PsnB | [1] |
| pYH001 | pET28b-6His-[Tm]-PsnB (R192A) | This study |
| pYH003 | pET28b-6His-[Tm]-PsnB (R181A) | This study |
| pYH006 | pET28b-6His-[Tm]-PsnB (L196A) | This study |
| pYH007 | pET28b-6His-[Tm]-PsnB (V201A) | This study |
| pYH010 | pET28b-6His-[Tm]-PsnB (Y171A) | This study |
| pYH012 | pET28b-6His-[Tm]-PsnB (F203A) | This study |
| pYH013 | pET28b-6His-[Tm]-PsnB (R101A) | This study |
| pYH014 | pET28b-6His-[Tm]-PsnB (R213A) | This study |
| pIN035 | pET28b-6His-[Tm]-PsnB (I193A) | This study |
| pHB100 | pET28b-6His-[Tm]-DLFIEDLGKVT-$(GS)_{10}$-PsnB | [6] |
| pIN036 | pET28b-6His-[Tm]-DLFIEDLGKVT-$(GS)_{10}$-PsnB (R72A) | This study |
| pYH023 | pET28b-6His-[Tm]-DLFIEDLGKVT-$(GS)_{10}$-PsnB (R101A) | This study |
| pYH024 | pET28b-6His-[Tm]-DLFIEDLGKVT-$(GS)_{10}$-PsnB (R213A) | This study |
| pIN028 | pET28b-6His-[Tm]-PsnB (D232A) | This study |
| pIN029 | pET28b-6His-[Tm]-PsnB (F233A) | This study |
| pIN030 | pET28b-6His-[Tm]-PsnB (R234A) | This study |

**Supplementary table 3. Oligonucleotides for cloning in this study**

| Plasmids | Primers |
|---|---|
| pYH001 | 5'-CAGACAGACGTTCAATCGCATCAGCTTCCAGATCTTTCGC<br>5'-GGAAGCTGATGCGATTGAACGTCTGTCTGCAGC |
| pYH003 | 5'-GGTGCACGCACCGCCAAACTGGAAGCG<br>5'-CGCTTCCAGTTTGGCGGTGCGTGCACC |
| pYH006 | 5'-CAGGTGCTGCAGACGCACGTTCAATGCGATCAGCTTCC<br>5'-CATTGAACGTGCGTCTGCAGCACCTGTTTGTTTTCAGG |
| pYH007 | 5'-CAGTTCCTGAAAACACGCAGGTGCTGCAGACAGACG<br>5'-CGTCTGTCTGCAGCACCTGCGTGTTTTCAGGAACTGACCGGC |
| pYH010 | 5'-GTTGGCGATTGCATTGCGAAACCTGTTGCT<br>5'-AGCAACAGGTTTCGCAATGCAATCGCCAAC |
| pYH012 | 5'-GCACCTGTTTGTGCCCAGGAACTGCTG<br>5'-CAGCAGTTCCTGGGCACAAACAGGTGC |
| pYH013, 023 | 5'-CTGCTGGCCTTTGCCGAACGCAGCACCC<br>5'-GGGTGCTGCGTTCGGCAAAGGCCAGCAG |
| pYH014, 024 | 5'-CCGGCGATGATGTTGCGGTGTATGTTATTGATGATCAGG<br>5'-CCTGATCATCAATAACATACACCGCAACATCATCGCCGG |
| pIN035 | 5'-GCGGAACGTCTGTCTGCAGCACC<br>5'-GCGATCAGCTTCCAGATCTTTCGCTT |
| pIN036 | 5'-GCGAGTCTGTATCAGAGCCCGGGC<br>5'-CAGATACACGGCGGCCGGACG |
| pIN028 | 5'-GCGTTTCGCCAGGCGGAAGAACGCATTGAAGC<br>5'-GATTTCATCGGTAACAATGCGCAGAGCACA |
| pIN029 | 5'-GATGCACGTCAGGCGGAAGAACGCATTGAAGC<br>5'-GATTTCATCGGTAACAATGCGCAGAGCACA |
| pIN030 | 5'-GATTTTGCCCAGGCGGAAGAACGCATTGAAGC<br>5'-GATTTCATCGGTAACAATGCGCAGAGCACA |

**Supplementary table 4. Observed and calculated mass values of MALDI data.** Difference is given as the observed mass minus the calculated mass.

| Ion | Observed | Calculated | Difference |
|---|---|---|---|
| **0.1min, 0** | 2565.16 | 2565.34 | -0.18 |
| **7.5min, 2** | 2529.13 | 2529.3 | -0.17 |
| **7.5min, 1** | 2547.24 | 2547.32 | -0.08 |
| **15min, 2** | 2529.09 | 2529.3 | -0.21 |
| **15min, 1** | 2547.14 | 2547.32 | -0.18 |
| **30min, 2** | 2529.1 | 2529.3 | -0.2 |

**Supplementary Figure 1d**

| Ion | Observed | Calculated | Difference |
|---|---|---|---|
| **1-a** | 2564.97 | 2565.34 | -0.37 |
| **2-a** | 2546.91 | 2547.32 | -0.41 |
| **3-a** | 2528.87 | 2529.3 | -0.43 |
| **1-c** | 2579.9 | 2580.35 | -0.45 |
| **2-c** | 2561.95 | 2562.33 | -0.38 |

**Supplementary Figure 2b left (0.5 M HA)**

| Ion | Observed | Calculated | Difference |
|---|---|---|---|
| **3-a** | 2529.01 | 2529.3 | -0.29 |

**Supplementary Figure 2b right (0 M HA)**

| Ion | Observed | Calculated | Difference |
|---|---|---|---|
| **2-c** | 2562.02 | 2562.33 | -0.31 |
| **2-d** | 2594.04 | 2594.33 | -0.29 |

**Supplementary Figure 2c**

| Ion | Observed | Calculated | Difference |
|---|---|---|---|
| **b2** | 261.26 | 261.16 | 0.1 |
| **b5** | 618.43 | 618.31 | 0.12 |
| **b9** | 1015.71 | 1015.58 | 0.13 |
| **b10** | 1116.73 | 1116.63 | 0.1 |
| **b13** | 1358.77 | 1358.77 | 0 |
| **b14** | 1415.84 | 1415.79 | 0.05 |
| **b15** | 1472.93 | 1472.81 | 0.12 |
| **b17** | 1732.96 | 1732.93 | 0.03 |
| **b18** | 1833.98 | 1833.98 | 0 |
| **b19** | 1935.04 | 1935.02 | 0.02 |
| **b20** | 2048.13 | 2048.11 | 0.02 |
| **b21** | 2119.13 | 2119.14 | -0.01 |
| **b22** | 2232.31 | 2232.23 | 0.08 |
| **b23** | 2289.36 | 2289.25 | 0.11 |
| **b24** | 2433.38 | 2433.29 | 0.09 |
| **y17** | 1664.88 | 1664.84 | 0.04 |
| **y19** | 1849.98 | 1849.95 | 0.03 |
| **y20** | 1963.12 | 1963.04 | 0.08 |
| **y22** | 2207.01 | 2207.11 | -0.1 |
| **$[M+H]^+$** | 2579.9 | 2580.35 | -0.45 |

**Supplementary Figure 2d (1-c)**

| Ion | Observed | Calculated | Difference |
|---|---|---|---|
| **b2** | 261.08 | 261.16 | -0.08 |
| **b5** | 618.18 | 618.31 | -0.13 |
| **b9** | 1015.39 | 1015.58 | -0.19 |
| **b10** | 1116.36 | 1116.63 | -0.27 |
| **b13** | 1358.42 | 1358.77 | -0.35 |
| **b14** | 1415.4 | 1415.79 | -0.39 |
| **b15** | 1472.4 | 1472.81 | -0.41 |
| **b17** | 1732.42 | 1732.93 | -0.51 |
| **b18** | 1833.56 | 1833.97 | -0.41 |
| **b19** | 1916.59 | 1917.02 | -0.43 |
| **b24** | 2399.9 | 2400.29 | -0.39 |
| **b25** | 2528.96 | 2529.33 | -0.37 |
| **y13** | 1332.31 | 1332.68 | -0.37 |
| **y14** | 1389.35 | 1389.7 | -0.35 |
| **y15** | 1446.35 | 1446.72 | -0.37 |
| **y16** | 1547.36 | 1547.77 | -0.41 |
| **y17** | 1646.36 | 1646.84 | -0.48 |
| **y18** | 1774.53 | 1774.93 | -0.4 |
| **y20** | 1944.59 | 1945.04 | -0.45 |
| **y22** | 2188.75 | 2189.11 | -0.36 |
| **y23** | 2301.88 | 2302.19 | -0.31 |
| **$[M+H]^+$** | 2562.02 | 2562.33 | -0.31 |

**Supplementary Figure 2d (2-c)**

| Ion | Observed | Calculated | Difference |
|---|---|---|---|
| **b2** | 261.26 | 261.16 | 0.1 |
| **b5** | 618.47 | 618.31 | 0.16 |
| **b9** | 1015.8 | 1015.58 | 0.22 |
| **b10** | 1116.87 | 1116.63 | 0.24 |
| **b13** | 1358.88 | 1358.77 | 0.11 |
| **b14** | 1415.9 | 1415.79 | 0.11 |
| **b15** | 1472.93 | 1472.81 | 0.12 |
| **b17** | 1732.95 | 1732.93 | 0.02 |
| **b18** | 1834.13 | 1833.98 | 0.15 |
| **b19** | 1935.12 | 1935.02 | 0.1 |
| **b20** | 2048.27 | 2048.11 | 0.16 |
| **b21** | 2119.27 | 2119.14 | 0.13 |
| **b22** | 2232.31 | 2232.23 | 0.08 |
| **b23** | 2289.31 | 2289.25 | 0.06 |
| **b24** | 2432.38 | 2432.29 | 0.09 |
| **b25** | 2561.49 | 2561.33 | 0.16 |
| **y16** | 1579.88 | 1579.77 | 0.11 |
| **y17** | 1678.88 | 1678.84 | 0.04 |
| **y19** | 1864.12 | 1863.95 | 0.17 |
| **y20** | 1977.14 | 1977.04 | 0.1 |
| **y21** | 2092.16 | 2092.07 | 0.09 |
| **$[M+H]^+$** | 2594.04 | 2594.33 | -0.29 |

**Supplementary Figure 2d (2-d)**

| Ion | Observed | Calculated | Difference |
|---|---|---|---|
| **PsnB,0** | 1150.39 | 1150.56 | -0.17 |
| **LP_PsnB,0** | 1150.42 | 1150.56 | -0.14 |
| **LP_PsnB,1** | 1132.42 | 1132.54 | -0.12 |
| **LP_PsnB,2** | 1114.41 | 1114.52 | -0.11 |

**Figure 2c**

| Ion | Observed | Calculated | Difference |
|---|---|---|---|
| **WT,0.1,0** | 2565.35 | 2565.34 | 0.01 |
| **WT,7.5,0** | 2565.39 | 2565.34 | 0.05 |
| **WT,7.5,1** | 2547.39 | 2547.32 | 0.07 |
| **WT,7.5,2** | 2529.33 | 2529.3 | 0.03 |
| **WT,15,2** | 2529.47 | 2529.3 | 0.17 |
| **T31A,0.1,0** | 2535.39 | 2535.33 | 0.06 |
| **T31A,7.5,0** | 2535.46 | 2535.33 | 0.13 |
| **T31A,15,0** | 2535.53 | 2535.33 | 0.2 |
| **T31A,30,0** | 2535.49 | 2535.33 | 0.16 |
| **T31A,30,1** | 2517.52 | 2517.31 | 0.21 |
| **T31A,60,0** | 2535.55 | 2535.33 | 0.22 |
| **T31A,60,1** | 2517.56 | 2517.31 | 0.25 |
| **T31A,120,1** | 2517.59 | 2517.31 | 0.28 |
| **T32A,0.1,0** | 2535.51 | 2535.33 | 0.18 |
| **T32A,7.5,0** | 2535.6 | 2535.33 | 0.27 |
| **T32A,15,0** | 2535.6 | 2535.33 | 0.27 |
| **T32A,30,0** | 2535.6 | 2535.33 | 0.27 |
| **T32A,60,0** | 2535.67 | 2535.33 | 0.34 |
| **T32A,120,0** | 2535.67 | 2535.33 | 0.34 |
| **31A32A,0.1,0** | 2505.53 | 2505.32 | 0.21 |
| **31A32A,7.5,0** | 2505.6 | 2505.32 | 0.28 |
| **31A32A,15,0** | 2505.64 | 2505.32 | 0.32 |
| **31A32A,30,0** | 2505.66 | 2505.32 | 0.34 |
| **31A32A,60,0** | 2505.7 | 2505.32 | 0.38 |
| **31A32A,120,0** | 2505.73 | 2505.32 | 0.41 |
| **E37A,0.1,0** | 2507.58 | 2507.33 | 0.25 |
| **E37A,7.5,0** | 2507.64 | 2507.33 | 0.31 |
| **E37A,7.5,1** | 2489.62 | 2489.31 | 0.31 |
| **E37A,15,0** | 2507.6 | 2507.33 | 0.27 |
| **E37A,15,1** | 2489.62 | 2489.31 | 0.31 |
| **E37A,30,0** | 2507.69 | 2507.33 | 0.36 |
| **E37A,30,1** | 2489.62 | 2489.31 | 0.31 |
| **E37A,60,1** | 2489.75 | 2489.31 | 0.44 |
| **E37A,120,1** | 2489.8 | 2489.31 | 0.49 |
| **E38A,0.1,0** | 2507.51 | 2507.33 | 0.18 |
| **E38A,7.5,0** | 2507.58 | 2507.33 | 0.25 |
| **E38A,7.5,1** | 2489.5 | 2489.31 | 0.19 |
| **E38A,15,0** | 2507.6 | 2507.33 | 0.27 |
| **E38A,15,1** | 2489.59 | 2489.31 | 0.28 |
| **E38A,30,0** | 2507.71 | 2507.33 | 0.38 |
| **E38A,30,1** | 2489.7 | 2489.31 | 0.39 |
| **E38A,60,0** | 2507.67 | 2507.33 | 0.34 |
| **E38A,60,1** | 2489.66 | 2489.31 | 0.35 |
| **E38A,120,1** | 2489.75 | 2489.31 | 0.44 |
| **37A38A,0.1,0** | 2449.54 | 2449.33 | 0.21 |
| **37A38A,7.5,0** | 2449.63 | 2449.33 | 0.3 |
| **37A38A,15,0** | 2449.67 | 2449.33 | 0.34 |
| **37A38A,30,0** | 2449.67 | 2449.33 | 0.34 |
| **37A38A,60,0** | 2449.72 | 2449.33 | 0.39 |
| **37A38A,120,0** | 2449.77 | 2449.33 | 0.44 |

**Supplementary Figure 4b**

| Ion | Observed | Calculated | Difference |
|---|---|---|---|
| **DFR,0.5,1** | 2547.89 | 2547.32 | 0.57 |
| **DFR,0.5,2** | 2529.84 | 2529.3 | 0.54 |
| **DFR,5,2** | 2529.84 | 2529.3 | 0.54 |
| **AFR,5,2** | 2565.99 | 2565.34 | 0.65 |
| **DAR,5,2** | 2565.99 | 2565.34 | 0.65 |
| **DFA,5,2** | 2565.88 | 2565.34 | 0.54 |
| **AAA,5,2** | 2565.99 | 2565.34 | 0.65 |

**Figure 4f**

| Ion | Observed | Calculated | Difference |
|---|---|---|---|
| **WT,0.1,0** | 2565.31 | 2565.34 | -0.03 |
| **WT,0.5,2** | 2529.08 | 2529.3 | -0.22 |
| **WT,32,2** | 2529.19 | 2529.3 | -0.11 |
| **R101A,0.1,0** | 2565.56 | 2565.34 | 0.22 |
| **R101A,0.5,2** | 2529.7 | 2529.3 | 0.4 |
| **R101A,32,2** | 2529.74 | 2529.3 | 0.44 |
| **R213A,0.1,0** | 2565.56 | 2565.34 | 0.22 |
| **R213A,0.5,0** | 2565.49 | 2565.34 | 0.15 |
| **R101A,32,0** | 2565.65 | 2565.34 | 0.31 |
| **R101A,32,1** | 2547.64 | 2547.32 | 0.32 |

**Figure 5f**

| Ion | Observed | Calculated | Difference |
|---|---|---|---|
| **WT,0.01,0** | 2565.31 | 2565.34 | -0.03 |
| **WT,0.5,2** | 2529.08 | 2529.3 | -0.22 |
| **WT,2,2** | 2529.24 | 2529.3 | -0.06 |
| **WT,8,2** | 2529.24 | 2529.3 | -0.06 |
| **WT,32,2** | 2529.19 | 2529.3 | -0.11 |
| **Y171A,0.01,0** | 2565.16 | 2565.34 | -0.18 |
| **Y171A,0.5,0** | 2565.26 | 2565.34 | -0.08 |
| **Y171A,0.5,1** | 2547.21 | 2547.32 | -0.11 |
| **Y171A,2,0** | 2565.24 | 2565.34 | -0.1 |
| **Y171A,2,1** | 2547.27 | 2547.32 | -0.05 |
| **Y171A,2,2** | 2529.24 | 2529.3 | -0.06 |
| **Y171A,8,1** | 2547.25 | 2547.32 | -0.07 |
| **Y171A,8,2** | 2529.24 | 2529.3 | -0.06 |
| **Y171A,32,1** | 2547.23 | 2547.32 | -0.09 |
| **Y171A,32,2** | 2529.21 | 2529.3 | -0.09 |
| **R181A,0.01,0** | 2565.4 | 2565.34 | 0.06 |
| **R181A,0.5,0** | 2565.31 | 2565.34 | -0.03 |
| **R181A,0.5,1** | 2547.37 | 2547.32 | 0.05 |

| | | | |
|---|---|---|---|
| **R181A,0.5,2** | 2529.31 | 2529.3 | 0.01 |
| **R181A,2,2** | 2529.35 | 2529.3 | 0.05 |
| **R181A,8,2** | 2529.31 | 2529.3 | 0.01 |
| **R181A,32,2** | 2529.47 | 2529.3 | 0.17 |
| **R192A,0.01,0** | 2565.38 | 2565.34 | 0.04 |
| **R192A,0.5,0** | 2565.4 | 2565.34 | 0.06 |
| **R192A,0.5,1** | 2547.34 | 2547.32 | 0.02 |
| **R192A,0.5,2** | 2529.26 | 2529.3 | -0.04 |
| **R192A,2,2** | 2547.3 | 2547.32 | -0.02 |
| **R192A,8,2** | 2529.28 | 2529.3 | -0.02 |
| **R192A,32,2** | 2529.51 | 2529.3 | 0.21 |
| **I193A,0.01,0** | 2529.4 | 2529.3 | 0.1 |
| **I193A,0.5,0** | 2565.38 | 2565.34 | 0.04 |
| **I193A,0.5,1** | 2565.47 | 2565.34 | 0.13 |
| **I193A,0.5,2** | 2547.41 | 2547.32 | 0.09 |
| **I193A,2,2** | 2529.42 | 2529.3 | 0.12 |
| **I193A,8,2** | 2529.51 | 2529.3 | 0.21 |
| **I193A,32,2** | 2529.4 | 2529.3 | 0.1 |
| **L196A,0.01,0** | 2529.54 | 2529.3 | 0.24 |
| **L196A,0.5,0** | 2565.38 | 2565.34 | 0.04 |
| **L196A,2,0** | 2565.49 | 2565.34 | 0.15 |
| **L196A,8,0** | 2565.49 | 2565.34 | 0.15 |
| **L196A,32,0** | 2565.51 | 2565.34 | 0.17 |
| **V201A,0.01,0** | 2565.51 | 2565.34 | 0.17 |
| **V201A,0.5,0** | 2565.56 | 2565.34 | 0.22 |
| **V201A,0.5,1** | 2565.49 | 2565.34 | 0.15 |
| **V201A,0.5,2** | 2547.44 | 2547.32 | 0.12 |
| **V201A,2,2** | 2529.42 | 2529.3 | 0.12 |
| **V201A,8,2** | 2529.49 | 2529.3 | 0.19 |
| **V201A,32,2** | 2529.58 | 2529.3 | 0.28 |
| **F203A,0.01,0** | 2529.58 | 2529.3 | 0.28 |
| **F203A,0.5,0** | 2565.51 | 2565.34 | 0.17 |
| **F203A,2,0** | 2565.47 | 2565.34 | 0.13 |
| **F203A,8,0** | 2565.61 | 2565.34 | 0.27 |
| **F203A,32,0** | 2565.68 | 2565.34 | 0.34 |

**Supplementary Figure 12e**

### Supplementary Note

#### Chemistry

#### 1. General Information

Unless otherwise noted, all reactions were performed under inert conditions. Nuclear magnetic resonance (NMR) spectra were recorded in $CDCl_3$ or $CD_3CN$ on Varian 400 NMR (400 MHz) spectrometers, with the residual solvent signal was used as a reference. High-resolution mass spectrometry (HRMS) was performed at the Organic Chemistry Research Center in Sogang University or at the Department of Chemistry in Seoul National University using the electrospray ionization (ESI) method. Chemical shifts are reported in ppm and coupling constants are given in Hz. Reactions were monitored by thin-layer chromatography (TLC) on EMD Silica Gel 60 F254 plates, and visualized either using UV light (254 nm) or by staining with potassium permanganate and heating. Dichloromethane ($CH_2Cl_2$) and tetrahydrofuran (THF) were dried using a PureSolv solvent purification system. Deuterated compounds were purchased from Cambridge Isotope Laboratories, Inc. and Sigma-Aldrich Corporation.

#### 2. Reagents

Diethyl (difluoromethyl)phosphonate (Sigma), Bromotrimethylsilane (Acros), Oxalyl chloride (Alfa), Dichloromethane (Acros), N,N-diisopropylethylamine (Alfa), Benzyl alcohol (Samchun), Pyridine (Samchun), Tetrahydrofuran (Fisher), Fmoc-Glu-OH (GL biochem), Acetic anhydride (Samchun), Diisopropylamine (Acros), n-Butyllithium (Acros), Ammonium chloride (Samchun).

#### 3. Substrate Preparation

**Dibenzyl (difluoromethyl)phosphonate (4)**

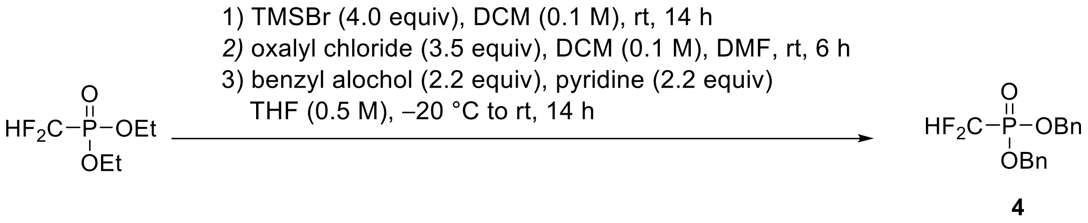


Dibenzyl (difluoromethyl)phosphonate was synthesized via a previously reported procedure with slight modifications[2]. To a stirred solution of diethyl (difluoromethyl)phosphonate (1.50 g, 8.0 mmol) in anhydrous

$CH_2Cl_2$ (8 mL) was added bromotrimethylsilane (4.90 g, 32.0 mmol) at room temperature. The reaction mixture was stirred for 14 h at the same temperature and then concentrated to give a residue. To a stirred solution of this residue in anhydrous $CH_2Cl_2$ (8 mL) was slowly added oxalyl chloride (3.55 g, 28.0 mmol) and DMF (a few drops) at 0 ºC. After being stirred for 6 h at room temperature, the reaction mixture was concentrated to give a residue. To a stirred solution of this residue in THF (16 mL) was added benzyl alcohol (1.90 g, 17.6 mmol) and pyridine (1.39 g, 17.6 mmol) at 20 ºC. After the resulting mixture was stirred for 30 min at the same temperature, stirring was continued for 6 h at room temperature before it was quenched with $KHSO_4$ (sat. aq, 15 mL). The layers were separated and the aqueous layer was extracted with EtOAc (3 × 15 mL). The combined organic extracts were dried (anhydrous $Na_2SO_4$) and concentrated in *vacuo*. The resulting residue was purified by flash column chromatography (silica gel, hexanes/ EtOAc, gradient elution) to afford **4** (730 mg, 2.7 mmol, 29% yield). All physical and spectroscopic data were in accordance with the literature[7].

**(9H-Fluoren-9-yl)methyl (S)-(2,6-dioxotetrahydro-2H-pyran-3-yl)carbamate (5)**

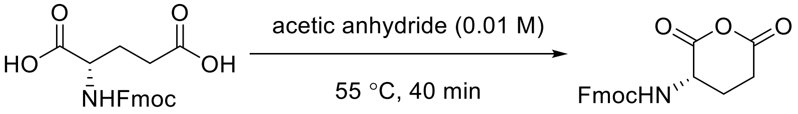


**5**

(((9H-fluoren-9-yl)methoxy)carbonyl)glutamic acid (738.7 mg, 2.0 mmol) and acetic anhydride (200 mL) were heated to 55 ºC until the solution turned clear and then were stirred for an additional 40 min at 55 ºC. The solution was cooled and evaporated under reduced pressure (12 mbar) at 55 ºC. The resulting solid was dried under vacuum to afford **5** (604 mg, 1.72 mmol, 86% yield).

$^1$H NMR (400 MHz, $CD_3CN$): δ = 7.84 (d, *J* = 7.5 Hz, 2H), 7.68 (d, *J* = 6.9 Hz, 2H), 7.43 (t, *J* = 7.4 Hz, 2H), 7.35 (t, *J* = 7.3 Hz, 2H), 6.33 – 6.15 (m, 1H), 4.54 – 4.44 (m, 1H), 4.42 – 4.34 (m, 2H), 4.26 (t, *J* = 6.7 Hz, 1H), 2.96 – 2.83 (m, 2H), 2.14 – 2.05 (m, 2H); $^{13}$C NMR (101 MHz, $CD_3CN$): δ = 168.35, 167.95, 167.36, 144.98, 142.10, 128.69, 128.09, 126.10, 120.96, 67.55, 51.42, 47.89, 30.51, 23.36; HRMS-ESI (m/z) $[M+Na]^+$ calcd for $C_{20}H_{17}NO_5Na$, 374.1004; found: 374.0990

**(2S)-2-((((9H-fluoren-9-yl)methoxy)carbonyl)amino)-6-((benzyloxy)(hydroxy)phosphoryl)-6,6-difluoro-5-oxohexanoic acid (6)**

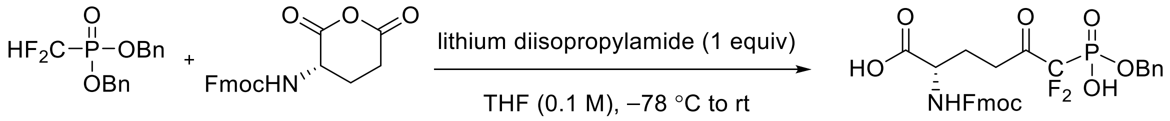


4 5 6

To a stirred solution of diisopropylamine (50.6 mg, 0.5 mmol) in anhydrous THF (2.0 mL) was added a solution of *n*-BuLi (312.5 μL, 1.6 M, 0.5 mmol) in anhydrous THF dropwise at –40 °C. The reaction mixture was warmed up to 0 °C, stirred for 30 min, and cooled down again to –78 °C. To the reaction mixture was added **4** (156.1 mg, 0.5 mmol) in anhydrous THF (0.5 mL) dropwise for 10 min at –78 ºC. After 1 h, the corresponding anhydride **5** (184.7 mg, 0.5 mmol) in anhydrous THF (2.5 mL) was added dropwise to the reaction mixture at –78 ºC. After Stirring for 1 h at –78 ºC, stirring was continued for 10 h at room temperature before it was quenched with $NH_4Cl$ (sat. aq, 5 mL). The layers were separated and the aqueous layer was extracted with EtOAc (5 × 5 mL). The combined organic extracts were dried (anhydrous $Na_2SO_4$) and concentrated in *vacuo*. The resulting residue was purified by preparative reversed phase HPLC chromatography (Agilent 1260 Infinity / ZORBAX SB-C18 Semi-preparative column (9.4 x 250 mm, 5 μm particle size; Agilent)) to afford **6** (47 mg, 0.082 mmol, 16% yield, elution time: 11.08 min at linear gradient elution program from 58% to 64% eluent B for 13 min (0-2 min: 40% eluent B, 2-15 min: gradient) at a 4.0 ml/min flow rate) which was used in the next reaction.

$^1$H NMR (400 MHz, $CDCl_3$) δ 7.79 – 7.71 (m, 2H), 7.61 – 7.53 (m, 2H), 7.42 – 7.27 (m, 9H), 5.54 – 5.47 (m, 1H), 5.13 (s, 2H), 4.48 – 4.35 (m, 3H), 4.23 – 4.17 (m, 1H), 2.63 – 2.43 (m, 2H), 2.35 – 2.24 (m, 1H), 2.12 – 2.00 (m, 1H) ppm. $^{13}$C NMR (101 MHz, $CDCl_3$) δ 175.28, 173.10, 156.35, 143.74, 141.45, 135.70, 128.75, 128.52, 128.46, 127.90, 127.24, 125.20, 120.14, 67.37, 66.89, 53.36, 47.27, 30.51, 29.85, 27.42 ppm*; $^{19}$F NMR (376 MHz, $CDCl_3$) δ –114.38 (d, J = 107.1 Hz) ppm**; HRMS-ESI (m/z) $[M-H]^-$ calcd for $C_{28}H_{25}F_2NO_8P$, 572.1286; found: 572.1291

* The $^{13}$C signals corresponding to the ketone carbon and the fluorinated alpha carbon could not be detected due to the weak signals originating from the 3J and 2J C-F coupling, respectively[8].

** The $^{19}$F NMR chemical shifts for related compounds are reported[9, 10].
